# A transcription-driven microgel network poised near the sol-gel transition

**DOI:** 10.1101/2024.06.16.599208

**Authors:** Mattia Marenda, Michael Chiang, Rafal Czapiewski, Davide Michieletto, Jon Stocks, Sophie M Winterbourne, Jamilla Miles, Olivia CA Fleming, Elena Lazarova, Graeme R Grimes, Hannes Becher, Atlanta Cook, Ryu-Suke Nozawa, Davide Marenduzzo, Nick Gilbert

**Author notes:** Corresponding authors: Nick Gilbert, Ryu-Suke Nozawa Davide Marenduzzo.

## Abstract

Living cells are organised through active, non-equilibrium processes, yet the physical principles governing their material states remain unclear. The cell nucleus has been described as either fluid-like or solid-like, but these behaviours have not been reconciled within a single framework. Here we show that nascent RNA and the multivalent RNA-binding protein SAF-A assemble into a nucleus-spanning microgel network maintained near the sol–gel transition. This active network exhibits scale- and time-dependent viscoelasticity and naturally reconciles solid-like and fluid-like responses. By integrating super-resolution imaging with mechanistic polymer modelling, we identify RNA production and degradation as control parameters that tune network connectivity. The model predicts enhanced susceptibility to modest perturbations near criticality, which we confirm experimentally by shifting the system between sol-like and gel-like regimes. Measurements of molecular transport, combined with simulations, reveal size-dependent diffusion and transient trapping within gel pores, establishing direct physical consequences of the near-critical state. Our results identify RNA metabolism as an energy-driven regulator of nuclear material properties and position the nucleus as a living example of an active microgel poised near criticality.

---

Living cells are organised by active, non-equilibrium processes whose physical principles remain incompletely understood. The nuclear interior is a striking example, it supports essential processes, such as transcription and replication, within a crowded microenvironment composed of chromatin, ribonucleoproteins, enzymes, and nucleotides^1–4^, yet its material nature is unclear. Classical models describe the nucleus as a solid like scaffold^5–8^, whereas live-cell imaging show that many nuclear proteins are highly mobile^9,10^, and fluorescent tagging fails to visualise stable nuclear structures, suggesting a fluid-like behaviour. A unified physical framework reconciling these observations is lacking^11^.

Many nuclear proteins belong to the heterogeneous nuclear ribonucleoprotein (hnRNP) family^12^, several of which self-associate in an RNA-dependent manner^13–15^, consistent with soft-matter assembly driven by multivalent interactions. One prominent hnRNP is SAF-A (scaffold attachment factor A, HNRNPU)^16,17^, which possesses structured domains, intrinsically disordered regions and RNA-binding motifs^18,19^, and plays key roles in chromatin architecture and transcription^16,20^. Disruption of SAF-A causes severe developmental and neurological phenotypes^21–25^, underscoring its importance, and its structural features make it a candidate component of emergent nuclear material states.

Chromatin, the complex between DNA and histone proteins, exhibits scale-dependent mechanics. FRAP experiments indicate that chromatin domains are relatively immobile^26^, whereas high-resolution imaging reveals dynamic^27^ liquid-like domains at smaller scales^28^. In vitro, chromatin can phase separate into liquid-like condensates^29^, highlighting the importance of multivalent interactions. Transcriptionally active chromatin regions coalesce into ∼90-nm hubs visualised by RNA polymerase II staining^30,31^, and enriched in newly synthesised RNA^32^, proposed to arise through bridging-induced phase separation driven by multivalent chromatin– protein interactions^33,34^. However, how such local microphase separation determines the global material state of the nucleus remains unresolved.

Here, using super-resolution imaging integrated with mechanistic polymer modelling we show that nascent RNA and SAF-A assemble into a nucleus-spanning network of weakly interconnected microgels (∼120 nm in diameter), maintained near the sol-gel transition. We identify RNA production and degradation as parameters governing network connectivity, that operates close to criticality, such that modest perturbations in transcription or RNA turnover shift it between sol-like and gel-like regimes. This near-critical organisation naturally reconciles solid-like and fluid-like responses through scale-and time-dependent viscoelasticity.

We further demonstrate that this active microgel state has measurable physical consequences: it modulates intranuclear transport in a size-dependent manner, altering effective diffusivity and enabling transient trapping within gel pores. These indicate that the nucleus is a living example of a soft-matter system where material properties are tuned by continuous biochemical energy input. By linking RNA metabolism to the sol-gel transition we provide a quantitative physical framework for understanding nuclear accessibility, gene regulation, and the potential impact of disease-associated perturbations.

## SAF-A and newly synthesised RNA form interacting clusters

To explore intranuclear supramolecular structures under physiological conditions, we used super-resolution direct Stochastic Optical Reconstruction Microscopy (dSTORM) in RPE1 cells to examine the nuclear localisation of SAF-A and newly transcribed RNA (Fig. 1; Methods). SAF-A exhibited a punctate, clustered distribution, consistent with our previous observations^35^ (Fig. 1A). Clustering analysis using the Ripley H function, an analysis method used to characterise spatial point patterns over a given area, revealed a peak at ∼40 nm, corresponding to an estimated cluster diameter of ∼100 nm (calculated assuming clusters are spherical) (Fig. 1B). From cluster density measurements and nuclear volume scaling, we estimated ∼3700 ± 1500 SAF-A clusters per nucleus.

**Fig. 1.**
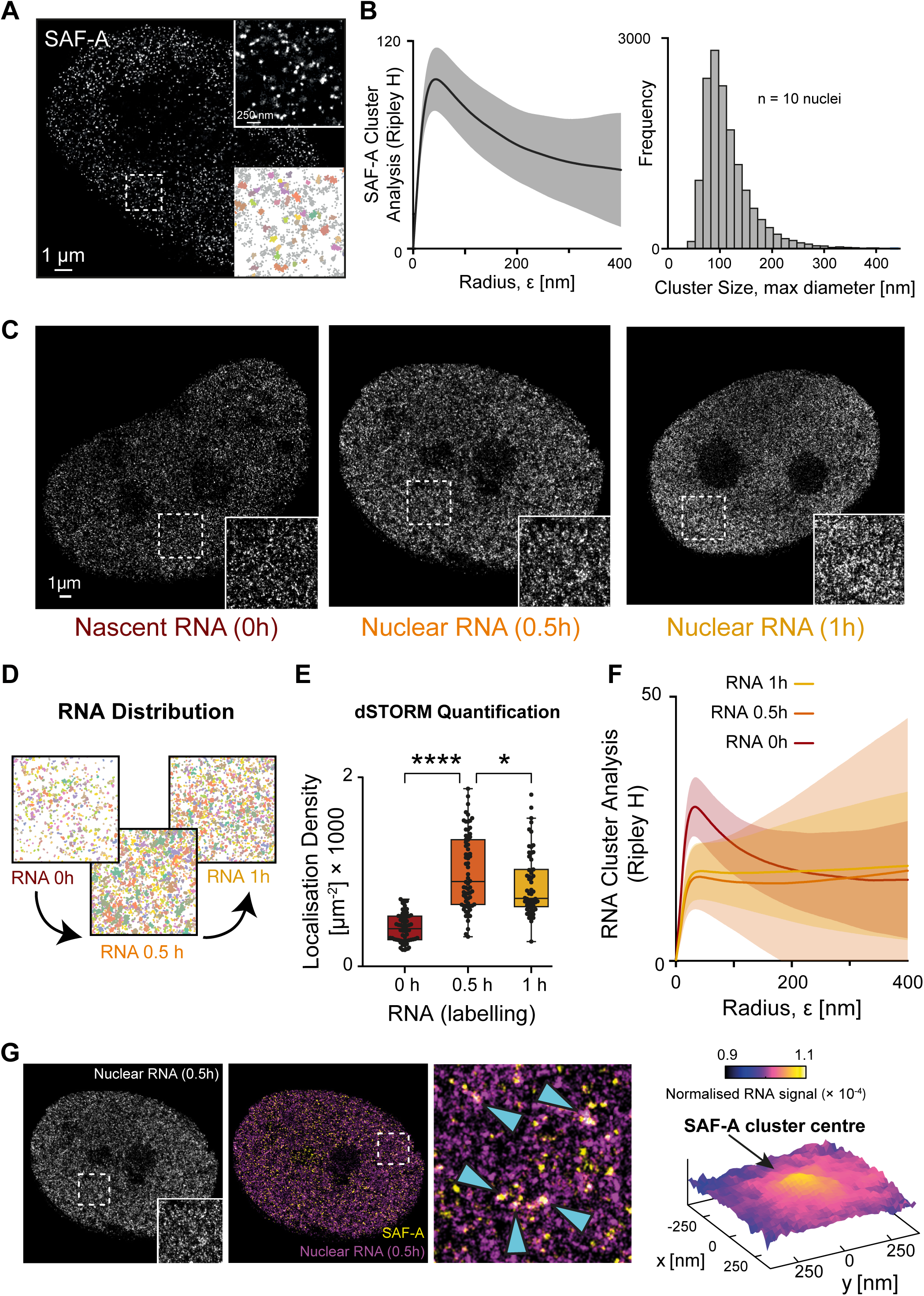
Organisation of SAF-A and nuclear RNA clusters. (**A**) STORM imaging of SAF-A clusters in fixed RPE1 cells. The top inset shows an enlarged view of region shown in dashed white square, whereas the bottom inset shows an X-Y plot of STORM localisations, with grey points not belonging to any cluster and coloured ones belonging to a cluster. (**B**) (Left) Strength of SAF-A clusters determined using Ripley H algorithm (n = 10 nuclei). (Right) Distribution of cluster size (n = 10 nuclei), computed as maximum distance between any two points in the cluster. (**C**) STORM imaging of nascent RNA and matured nuclear RNA. Cells were pulse labelled with 5EU then fixed at 0, 0.5, and 1 h. RNA was detected by attaching to CF647 dye using click chemistry. Insets are enlarged view of regions marked in dashed white square. (**D**) X-Y plot of STORM localisations shown in panel C insets, generated from the SuperStructure analysis using a fixed search radius. Different colours represent different detected clusters. (**E**) Quantification of RNA localisation density (n = 60 regions). *p* values for Student’s *t*-test: *, *p* < 0.05; ****, *p* < 0.0001. (**F**) Strength of RNA clusters for each time point (0, 0.5, and 1 h) determined using Ripley H algorithm (n = 10 nuclei). (**G**) (Left) Two-colour STORM imaging of SAF-A (yellow, CF568 dye) and RNA (purple, CF647 dye), at 0.5 h after RNA labelling. The left image shows RNA, with the inset corresponding to an enlarged view of the region marked by dashed box. The middle image show RNA and SAF-A, and the dashed box corresponds to the enlarged view shown in the right image. Blue arrows correspond to regions of pronounced SAF-A and RNA overlap. (Right) Meta-analysis of RNA signal strength surrounding SAF-A clusters (n = 10 nuclei).

Given that SAF-A binds RNA^7,16^, we next examined the relationship between SAF-A clustering and RNA. Pulse labelling with [5-^3^H]uridine showed that ∼80% of newly synthesised non-ribosomal RNA remained nuclear after 1 h, predominantly as high-molecular weight species (Fig. S1, A to C). For spatial analysis, RNA was labelled with 5-ethynyluridine (5EU) and visualised using click-chemistry (Fig. S1, D and E). Short 5EU pulses (15 min) enabled detection of nascent ribonucleoprotein structures, which were analysed by confocal microscopy to quantify RNA abundance (Fig. S1, F and G) and by STORM to analyse the spatial distribution (Fig. 1C).

Nascent RNA formed fine clusters (0 h; Fig. 1C), with RNA density increasing over time (Fig. 1, D and E). Ripley H analysis revealed a clustering peak at ∼20–30 nm at 0 h, which was lost by 0.5 h and 1 h, indicating RNA diffusion (Fig. 1F). As SAF-A is an RNA binding protein, we speculated that SAF-A clusters might form at sites of newly transcribed RNA. Consistently two-colour dSTORM showed significant colocalization of SAF-A and nascent RNA at early time points (Fig. 1G), followed by diffusion of RNA away from SAF-A clusters at later times (Fig. S1H).

## SAF-A is an RNA bridging factor

To assess whether experimental observations support a model in which SAF-A functions as an RNA-bridging protein we analysed its biophysical properties. Sequence analysis (Fig. S2A) and AlphaFold modelling (Fig. S2B) identified two RNA-binding sites in SAF-A: the known RGG domain and a previously uncharacterised charged surface on a domain that we term SPRNK, consisting of the SPRYdomain and an adjacent segment that encompassed a PNK-like NTPase domain with a weak affinity for ATPγS (KD = 806 μM) (Fig. S2C). Purified SPRNK bound to homomeric oligomers of A, C, or U, indicating that it binds RNA with little sequence specificity (Fig. S2, D and E) but RNA binding was enhanced in the presence of ATP (Fig. S2F), suggesting that RNA bridging might be dependent on local nucleotide concentration.

To further test the RNA binding activity of the SPRNK domain, four mutations were predicted based on surface electrostatics and conservation analysis of the AlphaFold model that might constitute an RNA binding surface. Mutation of these sites from positively charged to negatively charged amino acids (Fig. S2F) significantly abrogated RNA binding (Fig. S2, G to I). Furthermore, electron microscopy of cross-linked FLAG–SAF-A revealed filamentous oligomers that were disrupted by RNase treatment (Fig. S2, J and K), supporting the concept of RNA-mediated bridging.

## SAF-A and RNA form an intranuclear microgel network

To quantify the mesoscale organisation of RNA/SAF-A clusters (Fig. 1), we analysed cluster connectivity using the SuperStructure algorithm^35^ (Fig. S3A). This method measures how the number of detected clusters decay as the clustering radius is increased, providing a quantitative proxy for inter-cluster connectivity. SAF-A clusters (Fig. 1A) exhibited moderate connectivity, characterised by an exponential decay rate of 0.07 nm^-1^, corresponding to an effective connection probability of ∼0.05 between neighbouring clusters (Fig. 2A and Fig. S3B). Newly synthesised RNA clusters (Fig. 1C) were initially less interconnected (decay rate 0.03 nm^-1^; Fig. 2A and Fig. S3C), but connectivity increased over 0.5–1 h, coincident with the loss of discrete RNA clustering (Fig. 1F and Fig. S1H), approaching the connectivity regime observed for SAF-A. These measurements suggest that RNA/SAF-A assemblies are not isolated condensates but components of a weakly connected network.

**Fig. 2.**
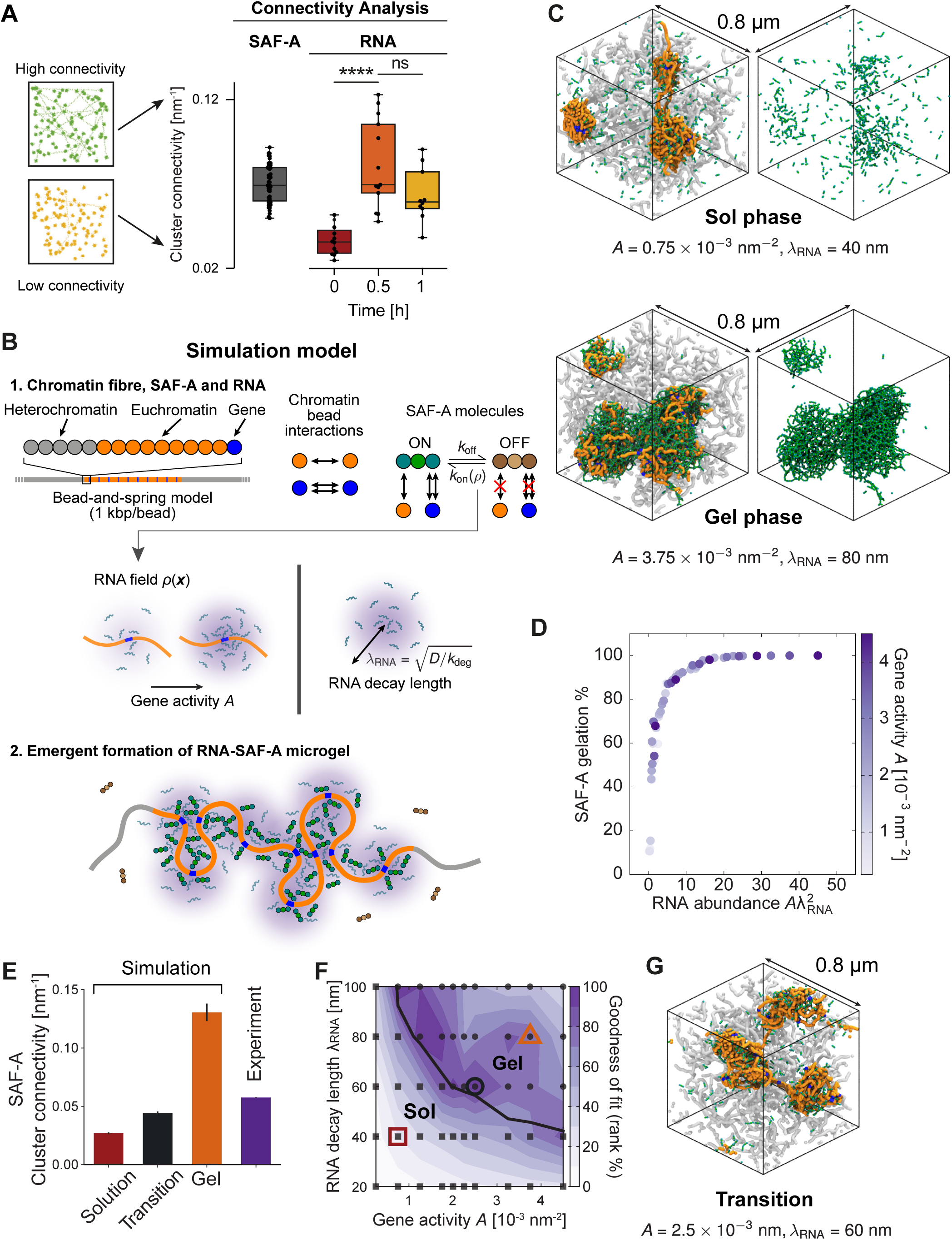
Simulations indicate that SAF-A and RNA form a network of microgels poised near the sol-gel transition. (**A**) (Left) Schematics depicting low and high cluster connectivity. (Right) Boxplots showing the extent of SAF-A cluster connectivity in RPE1 cells, compared to cluster connectivity in newly transcribed RNA, corresponding to data in Fig 1C. (**B**) Chromatin was modelled as a bead-and-spring copolymer made up by regularly interspaced heterochromatin (grey) and euchromatin (orange). Genes (blue) were interspersed within active chromatin. SAF-A molecules were modelled as patchy particles which can self-assemble to form oligomers when in the active phase (green). RNA molecules were modelled implicitly as a concentration field *ρ*(***r***), controlled by two parameters: gene activity *A* and RNA decay length λ_RNΑ_, which is dependent on RNA diffusion *D* and degradation *k*_deg_. The model can lead to gelation close to transcriptionally active regions of the chromatin fibre. (**C**) Snapshots of the whole system (left) and active SAF-A (right) in the sol phase (top) and gel phase (bottom); bead colours are the same as in (B). (**D**) Plot of fraction of active SAF-A proteins found in the largest cluster as a function of the combination *A*λ^2^ , showing that this is the main control parameter to determine when the sol-to-gel transition occurs within the parameter range analysed in simulations. (**E**) Bar chart showing cluster connectivity for simulations in the solution, transition, and gel regimes, compared to experimental data in (A). (**F**) Heatmap of the goodness of fit (rank of fit, in percentage from 0, worst, to 100, best) obtained by comparing simulated connectivity decay curves for active SAF-A to the experiments, as a function of *A* and λ_RNΑ_. The transition line (black) was plotted by analysing the gelation parameter introduced in (D). The best fits are close to the transition. (**G**) Simulation snapshot of the corresponding near-critical system [bead colours as in (B)].

In vitro, SAF-A and RNA form mesh-like assemblies^16^ (Fig. S2J), suggesting that the nuclear clusters correspond to hydrated RNA-protein condensates. Their self-limited size is characteristic of microphase separation^36^, consistent with RNA-rich condensates that assemble as microphase-separated hydrogels. Together, they can be summarised as microgels that form a sparse, nuclear-spanning network of weakly interconnected structures^37^. Such assemblies can arise through bridging-induced phase separation, a mechanism that generates microphase-separated clusters of transcription components, analagous to transcription factories^34,38^. While these structures can be visualised by super-resolution microscopy (Fig. 1, A and C), directly establishing their gel-like material properties remains challenging, as “microgels” describe a biophysical state rather than a discrete morphological entity.

To infer the physical state of the microgel network, we developed coarse-grained molecular dynamics simulations coupling polymer models of chromatin together with diffusing RNA and multivalent SAF-A particles (Fig. 2B, part 1; Methods). Chromatin was modelled as a semi-flexible polymer containing alternating transcriptionally active (euchromatin) and inactive (heterochromatin) regions, while SAF-A was represented as a multivalent particle which underwent self-association promoted by local RNA concentration. RNA production, diffusion, and degradation established by steady-state RNA concentration fields around active chromatin, which locally regulated SAF-A bridging, and led to the emergent formation of a RNA–SAF-A microgel (Fig. 2B, part 2) and network formation.

Increasing transcriptional output *A* or the RNA decay length λ_RNΑ_, which is dependent on RNA diffusion *D* and degradation *k*_deg_, drove a transition from a sol-like state, dominated by monomeric SAF-A (green), to a percolating gel-like network (Fig. 2C and movie S1 to S3). The degree of gelation depended primarily on total RNA abundance, scaling with the dimensionless parameter *A*λ^2^ (Fig. 2D). Comparison of experimental and simulated SuperStructure curves (Fig. 2E, see methods) revealed that neither fully sol-like nor fully gelled states recapitulated the approximately single-exponential decay observed in vivo. Instead, the closest agreement was obtained near the sol–gel transition (*A*λ^2^ ≈ 9; Fig. 2, E and F), creating a self-organised structure in cells, with weak interconnection between SAF-A–RNA clusters, poised near criticality (Fig. 2G)^39^.

## Intranuclear microgel network is dependent on SAF-A and transcription

If SAF-A bridges RNA to assemble an interconnected microgel network, then perturbing either SAF-A levels or transcription should disrupt network organisation (Fig. 3A). Consistent with this prediction, SAF-A depletion reduced the density and size of newly synthesised RNA clusters, indicative of increased RNA dispersion and weakened transcriptional hubs (Fig. 3, B and C). Inter-cluster connectivity was also reduced (Fig. 3C and Fig. S4), causing partial network collapse.

**Fig. 3.**
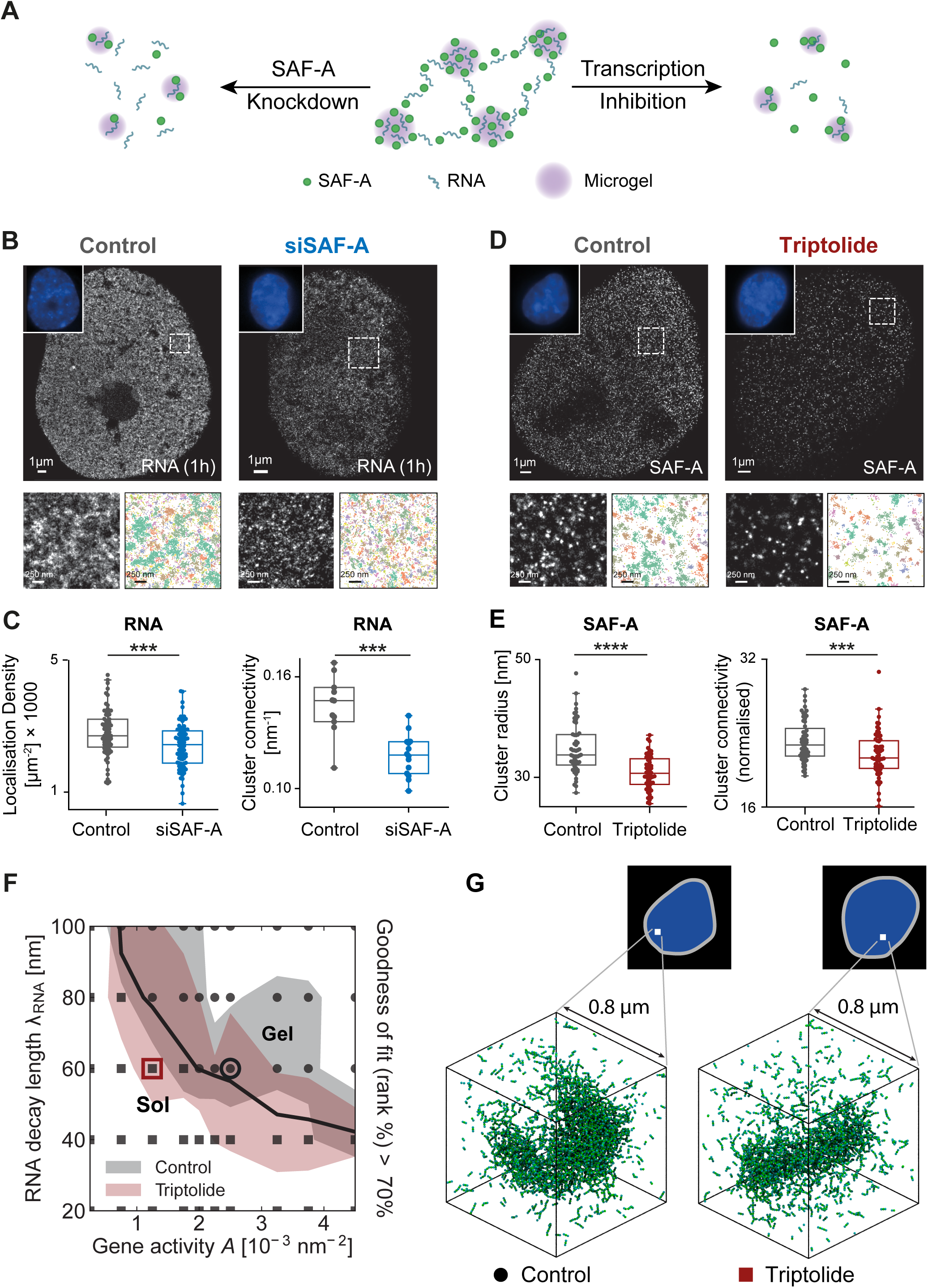
SAF-A depletion and transcriptional inhibition degrade microgel networks. (**A**) Cartoon outlining how a network of interconnected RNA–SAF-A microgels (middle) might be affected by SAF-A depletion (left) or transcription inhibition (right). (**B**) STORM of newly synthesised RNA labelled with CF647 dye in wild type (Control; left) and after SAF-A depletion (siSAF-A; right). DAPI staining is shown in the top left corner of each image. The lower left panel shows the enlarged view of regions marked in dashed white square, while the lower right panel is an X-Y plot of STORM localisations, generated from SuperStructure analysis for a fixed search radius. Colours represent different detected clusters. (**C**) Superstructure connectivity boxplots for siSAF-A cells, showing less connectivity compared to Control cells, indicating that SAF-A depletion disrupts the microgel network. *p* values for Mann-Whitney U test: ***, *p* < 0.001; ****, *p* < 0.0001. (**D**) Similar to (B), but showing STORM of SAF-A labelled with CF647 dye in wild-type cells (Control; left) and after transcription inhibition (Triptolide; right). (**E**) Superstructure connectivity plots after transcriptional inhibition by triptolide, showing a reduced connectivity with respect to the wild type (Control), suggesting that transcription inhibition disrupts the microgel network. *p* values for Mann-Whitney U test: ***, *p* < 0.001; ****, *p* < 0.0001 (nControl = 56 and nTriptolide = 64) (**F**) Phase diagram of simulated system with regions corresponding to the top 30% of fits between simulated and experimental connectivity decay curves for the wild type (grey, Control) and for transcription inhibition (pink, Triptolide). (**G**) Simulation snapshots within a 0.8 × 0.8 × 0.8 μm cube corresponding to parameter points within the best-fit region for each of the two conditions shown in (F): open circle for Control and open square for Triptolide.

We next inhibited transcription using triptolide or α-amanitin, both of which rapidly depleted nuclear RNA and produced similar phenotypes (Fig. S5, A and B). Triptolide treatment decreased SAF-A localisation and cluster density (Fig. 3D and Fig. S5C) and globally disrupted microgel connectivity (Fig. 3E), observed as a reduced decay rate in SuperStructure connectivity curves (Fig. S5D).

Quantitative comparison between experiments and simulations showed that the post-triptolide SuperStructure curves were best fit by smaller *A*λ^2^ values than wild-type, consistent with a shift toward the sol-like regime (Fig. 3F). Reduced RNA availability limits SAF-A bridging, increasing the fraction of inactive monomers and weakening network connectivity, thereby explaining the observed changes in microgel architecture (Fig. 3G).

## XRN2-mediated RNA turnover tunes microgel network morphology

Newly synthesised RNA is rapidly redistributed within the nucleus (Fig. 1C), suggesting that RNA turnover may regulate microgel organisation. We therefore used proximity-dependent biotin identification (BioID) to identify proteins spatially associated with SAF-A by tagging its N-terminus with BirA. In addition to known SAF-A interactors—including HNRNPUL1/2, SAFB, SAFB2, and MATRIN3 (Fig. S6A)—we unexpectedly identified an association to the 5′–3′ exonuclease XRN2.

XRN2 is a nuclear exonuclease involved in RNA degradation and transcription termination^8,40,41^. SAF-A and XRN2 interacted in an RNA-dependent manner (Fig. 4A), suggesting that XRN2 may degrade nascent RNA within SAF-A microgels (Fig. S6B). To test this, we measured nuclear RNA turnover following a 15-min 5EU pulse. In control cells, newly synthesised RNA accumulated transiently and decayed with a mean lifetime of ∼43 min (Fig. S7A). Depletion of XRN1 (Fig. S7B) or the exosome subunit EXOSC3 (Fig. S7C) had little effect on RNA persistence (Fig. S7A), whereas XRN2 knockdown (Fig. S7B) markedly stabilised nuclear RNA, increasing its lifetime to ∼140 min and promoting cytoplasmic accumulation at later time points (Fig. S7A). These findings were independently confirmed using an auxin-inducible degron allele of XRN2 (Fig. S7, D and E). Together, these results identify XRN2 as a mediator of nascent nuclear RNA turnover.

**Fig. 4.**
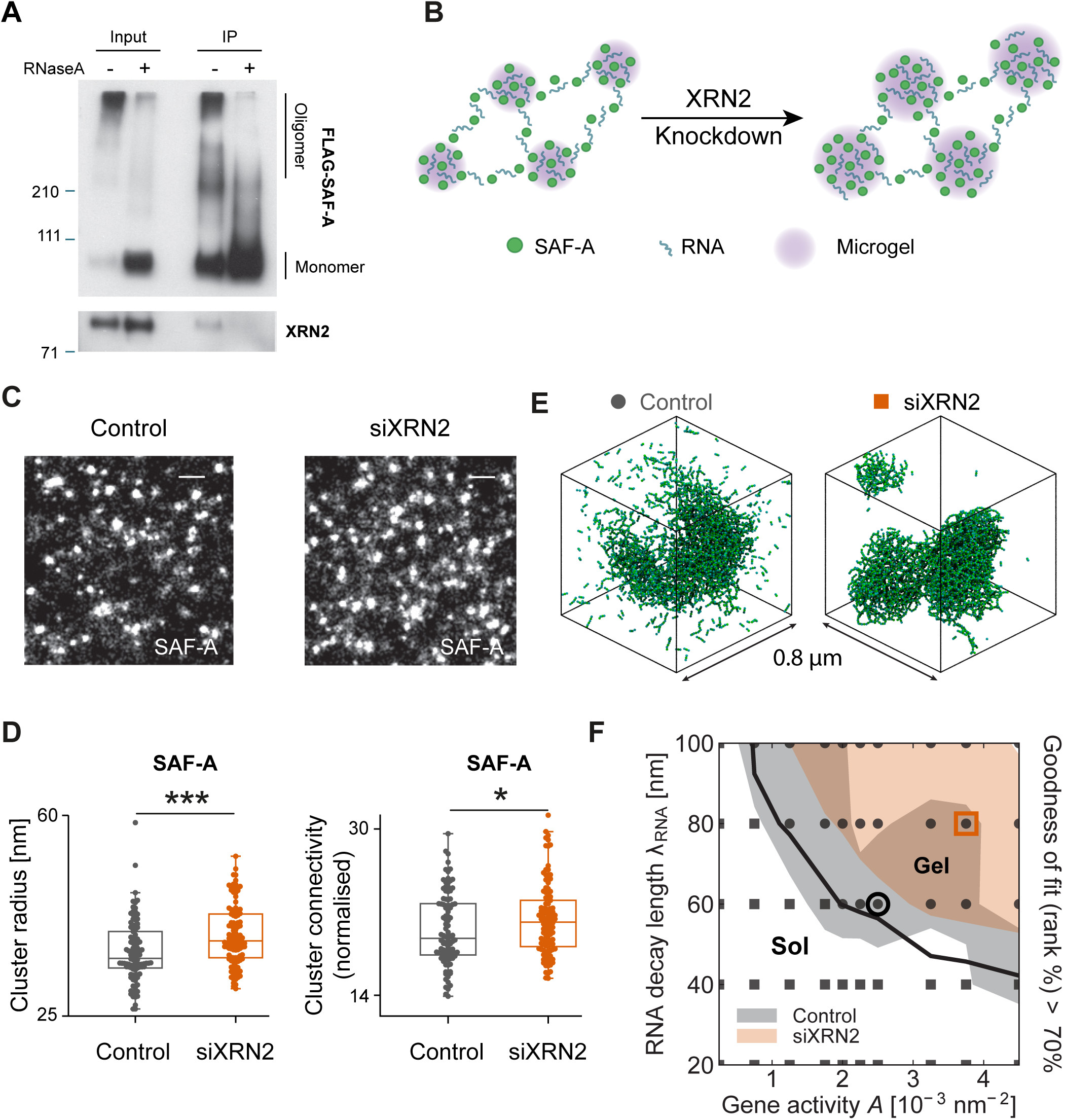
Effect of XRN2 on RNA–SAF-A microgel networks. (**A**) Western blots showing immunoprecipitation of oligomerised FLAG–SAF-A with XRN2, in the presence and absence of RNaseA. (**B**) Cartoon outlining how an interconnected network of RNA–SAF-A microgels (left) is influenced by XRN2 depletion (right). (**C**) STORM images of SAF-A in wild-type (Control) and XRN2-depleted (siXRN2) cells. Scale bar is 250 nm. (**D**) SAF-A cluster radius (left) and cluster connectivity (right) in Control and siXRN2 cells. *p* values for Mann-Whitney U test: ***, *p* < 0.001; ****, *p* < 0.0001 (nControl = 102 and nsiXRN2 = 112) (**E**) Simulation snapshots corresponding to parameter points within the best-fit regions shown in (F) when comparing connectivity decay curves between simulations and experiments for Control and siXRN2 conditions (open circle for Control and open square for siXRN2). (**F**) Phase diagram of simulated system with regions corresponding to the top 30% of fits between simulated and experimental connectivity decay curves for the wild-type (grey, Control) and XRN2 depletion cells (orange, siXRN2).

We next asked whether altered RNA turnover impacts microgel morphology (Fig. 4B). dSTORM imaging (Fig. 4C) revealed that XRN2 depletion increased SAF-A cluster size (Fig. 4D), consistent with an imbalance between RNA production and degradation. SuperStructure analysis showed increased inter-cluster connectivity following XRN2 knockdown (Fig. 4D and Fig. S8A), and comparison between experiments and simulations revealed best agreement at higher *A*λ^2^ values relative to wild-type (Fig. 4, E and F), indicative of a shift toward a more gel-like regime. Mechanistically, prolonged RNA residence near transcription sites provides additional substrate for SAF-A bridging, stabilising the network. Consistent with this interpretation, XRN2 depletion partially counteracted the effects of transcription inhibition by triptolide (Fig. S8, B and C), indicating that transcription and XRN2-mediated degradation act in opposition to dynamically tune microgel connectivity in the nucleus.

## Network morphology regulates intranuclear dynamics

Our results indicate that the intranuclear environment comprises a near-critical network of interconnected microgels poised close to the sol–gel transition. Perturbations such as SAF-A depletion, transcription inhibition, or XRN2 loss shift this balance toward sol-like or gel-like regimes. Because diffusion within a gel depends on pore size and connectivity, we hypothesised that changes in microgel morphology would directly affect intranuclear protein mobility, with potential consequences for transcriptional regulation.

To test this, we measured SAF-A dynamics using fluorescence correlation spectroscopy (FCS) and fluorescence recovery after photobleaching (FRAP). FCS revealed that transcription inhibition significantly increased SAF-A mobility (Fig. S9A). Consistently, FRAP analysis of SNAP–SAF-A also showed faster recovery following transcription inhibition (Fig. S9B), indicating microgel disruption, which was abrogated by loss of XRN2 (Fig. S9B).

To quantify how network morphology impacts diffusion more generally, we measured the mobility of GFP oligomers of defined sizes expressed from the ROSA26 locus (Fig. 5A and Fig. S9, C and D). As expected, larger oligomers diffused more slowly than monomers (7 vs. 22 μm^2^/s, respectively). Importantly, depletion of SAF-A increased the mobility of all GFP species (Fig. 5B), indicating that SAF-A contributes to an interconnected mesh that elevates local nuclear viscosity. Conversely, XRN2 depletion reduced GFP mobility (Fig. 5C), consistent with enhanced gelation.

**Fig. 5.**
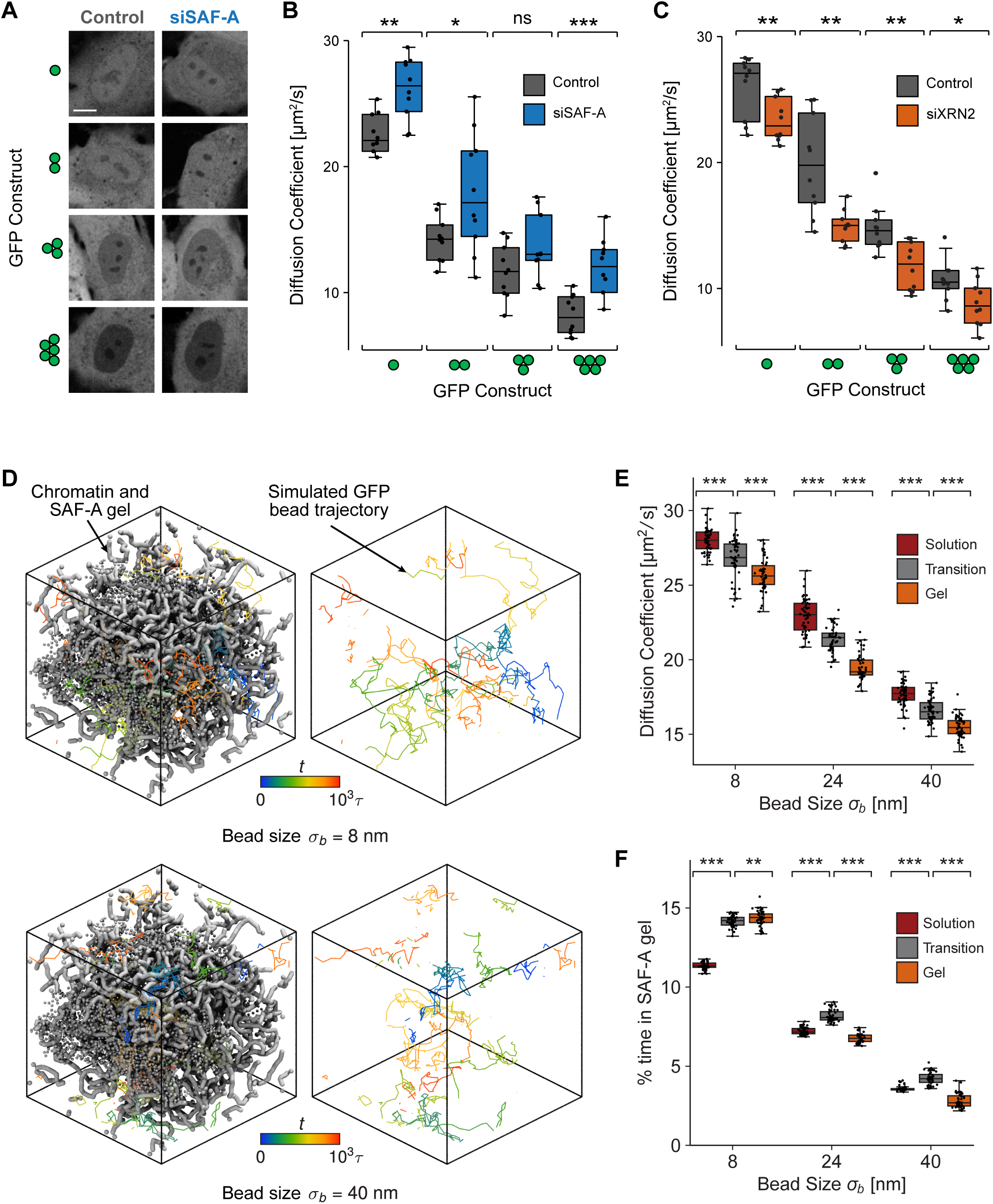
Impact of RNA–SAF-A microgel networks on protein mobility. (**A**) Live-cell imaging of RPE1 cells expressing mono, di, tri, and penta-GFP in wild-type cells (Control) and after SAF-A depletion (siSAF-A). Scale bar is 5 μm. (**B**) Boxplots of GFP diffusion coefficients, measured by FCS, in Control and siSAF-A cells expressing different GFP constructs (n = 10 regions). (**C**) Similar to (B), but for wild-type (Control) and XRN2 depleted (siXRN2) cells (n = 10 regions). (**D**) Trajectories of 8 nm (top) and 40 nm (bottom) spheres in the gel phase, corresponding to high transcriptional activity, showing that smaller particles are transiently trapped inside the gel, and larger ones are excluded from it. (**E**) Diffusion coefficient of different-size spheres (equivalent to GFP monomers and multimers) calculated from simulations in different conditions: sol phase, transition regime, and gel phase. (**F**). Boxplots showing the proportion of time a bead spends within the SAF-A gel (n = 50 simulations)*. p* values for Mann-Whitney U test: ns, not significant; *, *p* < 0.05; **, *p* < 0.01; ***, *p* < 0.001.

To better understand movement of particles in the nucleus, we complemented these experiments with simulations predicting the diffusion of particles of varying size in nuclear environments at different values of *A*λ^2^ , spanning sol-like, gel-like, and near-critical, or transition regimes (Fig. 5D). In all cases, effective diffusion coefficients *D*_eff_ were reduced relative to aqueous diffusion *D*_0_, as predicted by Stoke’s Law, and exhibited strong size dependence in the 8–40 nm range, reflecting confinement by the microgel pore structure. As diffusivity differed markedly between regimes, it supported our idea that nuclear viscosity is dynamically tunable (Fig. 5E).

Simulated trajectories further revealed dynamic heterogeneity: small particles were transiently trapped within the microgel network, whereas larger particles were excluded (Fig. 5F and movie S4 and S5). This behaviour is consistent with the “corral model” for intranuclear diffusion^42^, suggesting that the transcription-dependent microgel network acts as a size-selective sieve that can differentially sequester or exclude nuclear proteins.

Together, our experimental and computational results show that the near-critical RNA–SAF-A microgel network modulates intranuclear mobility and local viscosity in a size-and state-dependent manner. By tuning the balance between transcription and RNA degradation, cells can dynamically regulate protein accessibility and, consequently, gene expression.

## Discussion

We define the morphology and functional consequences of a dynamic intranuclear mesh formed by RNA and SAF-A. SAF-A assembles RNA into weakly interconnected, microphase-separated hydrogels (microgels) whose size and connectivity respond to transcriptional activity, RNA turnover, and SAF-A abundance. Because RNA mediates inter-microgel bridging, our findings reinforce the emerging view that RNA is a key structural determinant of nuclear organisation^7,43^. An important next step will be to examine how disease-associated SAF-A mutations—particularly within the RNA-binding SPRNK domain—alter microgel architecture and connectivity^3^.

Super-resolution imaging revealed ∼4,000 SAF-A microgels per nucleus, with an average diameter of ∼120 nm. Given estimates of ∼10,000 active genes in RPE1 cells, each microgel likely encompasses multiple regulatory elements and transcription units. The size and abundance of microgels closely match those reported for transcription factories^31^, suggesting a direct relationship. In this framework, transcription factories formed by bridging-induced phase separation^33^ may nucleate RNA–SAF-A microgels through local production of nascent RNA and SAF-A recruitment.

A key advance is the demonstration that microgels form a connected, nuclear-spanning network. Integrating dSTORM with simulations, we show that increasing transcriptional output or RNA decay length drives a transition from a sol-like regime, dominated by monomeric SAF-A, to a gel-like regime with a percolating RNA–SAF-A network. Quantitative comparison between experiment and theory indicates that, in vivo, this network operates near the sol–gel transition, consistent with a state of self-organised criticality. Such positioning enables high sensitivity to biological inputs, allowing small changes in transcription or RNA degradation to markedly tune network properties, as observed upon transcription inhibition or XRN2 depletion.

This near-critical organisation reconciles long-standing observations of both solid-like and fluid-like nuclear behaviour^26,28,44^ The RNA/SAF-A network constitutes a viscoelastic, heterogeneous environment whose material properties depend on length and time scale. Crucially, we show that continuous RNA degradation by XRN2 dynamically renews the network, preventing excessive gelation that would impede macromolecular diffusion. RNA turnover therefore maintains the nucleus in a functional regime that balances structural integrity with molecular mobility.

Functionally, microgel morphology directly regulates intranuclear dynamics. The network increases local viscosity near transcriptionally active regions in a size-dependent manner, selectively slowing or trapping proteins within the mesh. This provides a physical mechanism for concentrating transcription factors, co-activators, and RNA-processing machinery near active genes, while allowing rapid redistribution when required. In this way, the microgel network may contribute to the formation and regulation of transcriptional neighbourhoods ^45^

Finally, tunable gelation near highly active genes offers a potential biophysical explanation for transcription-induced chromatin decompaction observed in vivo^16,46–49^. Simulations suggest that strong local gelation can interfere with chromatin folding, leading to chromatin swelling and gene expansion. Further exploration of the material properties of RNA–SAF-A microgels—both in vitro and in cells—will be essential to fully understand how RNA dynamics shape nuclear mechanics and gene regulation.

## Acknowledgments

We thank members of the Gilbert and Marenduzzo labs for helpful discussions. We are grateful to Stuart Aitken (University of Edinburgh) for bioinformatics support, Jimi Wills (University of Edinburgh) for help with mass-spec, metabolomics and BioID2 analysis, the Institute of Genetics and Cancer Advanced Imaging Resource (AIR), and Kayoko Suenaga and Akihiko Ichikawa (Carl Zeiss Co. Ltd.) for supporting FCS analysis. Thanks to Steven West (University of Exeter) for supplying XRN2 degron HCT116 cells, Tomomi Kiyomitsu (OIST) for supplying the ROSA26 donor plasmid, and Karsten Rippe (German Cancer Research Center) for the multimeric GFP plasmids.

## Author contributions

Conceptualization: RSN, DMa, NG; Methodology: MM, MC, RC, DMi, JS, SMW, JM, OCAF, EL, GRG, HB, AC, RSN; Investigation: MM, MC, RC, DMi, JS, SMW, JM, OCAF, EL, GRG, HB, AC, RSN; Visualization: MM, MC, DMi; Funding acquisition: RSN, DMa, NG; Project administration: DMa, NG; Supervision: DMa, NG; Writing – original draft: MM, DMi, DMa, NG; Writing – review & editing: all

## Data and materials availability

Proteomics data are available via ProteomeXchange with identifier PXD053208. The following high-level data and analysis have been deposited in the Edinburgh DataShare (https://doi.org/10.7488/ds/7751): dSTORM, FRAP, FCS, and simulations. Simulation scripts and example initialization configurations which can be used to run the simulations using the LAMMPS software are available at https://git.ecdf.ed.ac.uk/chromatin-lab/nuclear_rna_gel. Scripts and raw data for the structural analysis of SAF-A (Fig. S2) are available at https://git.ecdf.ed.ac.uk/cooklab/hnrnpu. All raw data and further analysis scripts are available upon request. Data analysis and simulation software used are all open-source packages.

- All data are available in the manuscript or in the supplementary materials. All other data supporting the findings of this study are available from the lead contact upon request. All reagents generated in this study are available from the lead contact with a completed material transfer agreement if necessary
- This paper does not report original code.
- Any additional information required to reanalyze the data reported in this paper is available from the lead contact upon request.

## Materials and Methods

### Cell culture

hTERT-RPE1 (ATCC, cat# ATCC-CRL-4000), HCT116, 293T and FlpIn T-REx 293 cells were cultured as described previously^16,50,51^. Transcription was blocked by adding α-amanitin (50 µg.ml^-1^), triptolide (1 µM) and/or actinomycin D (50 ng.ml^-1^) to cells for the times indicated. To label newly transcribed RNA, cells were incubated with 5-ethynyl uridine (5EU, 0.5 mM), 4-thiouridine (4SU, 0.5 mM) or [5-^3^H]uridine for the indicated time. 5EU was fluorescently labelled with Alexa 647 or Alexa 488 through Click labelling as previously described ^16^. FlpIn T-Rex 293 stable cell line for doxycycline inducible expression of myc–BioID2–SAF-A was established as described^16^. hTERT-RPE1 stable cell lines for doxycycline inducible expression of GFP-multimers was established by electroporation with CRISPR-Cas9 and donor plasmids (see below for more details). Two days after electroporation, cells were selected with 2 mg.ml^-1^ neomycin and single clones were isolated. The cells were treated with 1 mg.ml^-1^ doxycycline for 24 hr for inducing expression of GFP oligomer proteins. To deplete XRN2 using the auxin-inducible degron system, auxin was added to cells at 500 nM for the indicated times.

### Plasmids

For the BioID2 assay, the myc-BioID2 fragment from the myc-BioID2-MCS plasmid (addgene #74223) was replaced with 3xFLAG in the FlpIn 3xFLAG–SAF-A plasmid^16^. For FRAP and FCS experiments, the SNAP gene was inserted between the 3xFLAG tag and SAF-A gene in the FlpIn 3xFLAG–SAF-A plasmid^16^. GFP-multimers fragment (gift from Karsten Rippe) were inserted into the Rosa26 donor plasmid (addgene #114699). For protein expression the SPRNK domain (amino acids 221-679) was synthesised as a G-block (IDT) and cloned into a T7 expression vector pEC_K_3C (gift from Atlanta Cook) by Gibson assembly. For the SPRNK RNA binding mutant the following mutations were introduced into the G-block: K246E, AAT-GAA; K251E, AAA-GAA; R539E, AGA-GAA; K616E, AAA-GAA. All plasmids were verified by DNA sequencing.

### Antibodies

anti-SAF-A 3G6, 1:5000 for western blotting, Abcam, ab10297. anti-SAF-A, 1:200 for immunofluorescence, Bethyl, A300-690A. anti-XRN1, 1:3000 for WB, 1:100 for IF, Bethyl, A300-443A anti-XRN2, 1:3000 for western blotting, 1:300 for immunostaining, Bethyl, A301-103A. anti-EXOSC3, 1:1000 for WB, 1:200 for IF, SantaCruz, sc98776 anti-GAPDH, 1:5000 for WB, Cell Signaling Technology, D16H11 anti-FLAG, 1:10000 for WB, Sigma, M2 anti-myc, 1:1000 for WB, Cell Signaling Technology, 2276S

### RNA interference

For siRNA treatment, cells (10%–20% confluent) were transfected with 10 nM Stealth RNA (ThermoFisher) using Lipofectamine RNAi MAX (ThermoFisher) for the time indicated. siRNA sequences and the identification numbers are CCUGGGAAUCGTGGCGGATATAATA HSS104917 for SAF-A, GCGAAAUGAUAACUUCUAATT s29016 for XRN1, GGAAAGUUGUGCAGUCGUATT s22412 for XRN2, UCAACUUUGAAUAUAUCUCCA s83102 for EXOSC3. The control RNA is Stealth RNAi siRNA Negative Control, Med GC (ThermoFisher).

### Domain analysis and alpha fold structure prediction

Sequence comparisons of SAF-A/HNRNPU with paralogues HNRNPUL1 and HNRNPUL2 were generated by selecting orthologous and paralogues sequences from model vertebrates using UNIPROT. Sequences were aligned using MAFFT with default settings^52^. The multiple sequence alignment was visualised using Jalview^53^. AlphaFold2 models of SAF-A (downloaded from AlphaFoldDB on 29/03/2023) were used for surface property analysis. Surface electrostatics were calculated using the adaptive Poisson-Boltzmann solver (APBS) plugin in PyMol with charge surfaces ranging from -3kT/e to +3kT/e^54^. Sequences for phylogenetic analysis of SAF-A were selected using the orthologous matrix database (OMA, group 116311)^55^ then supplemented with additional sequences by using PSI-BLAST. Conservation scores were calculated using ConSurf with the multiple sequence alignment and AlphaFold2 models as input^56^. Missense variants for the transcript ENST00000444376.2 were extracted from gnomAD (v2.1.1)^57^ and filtered for benign variants. Vd/Vp values were then calculated using the 1D-to-3D suite^58^. Protein conservation, missense variant data, secondary structure elements and phosphorylation sites were displayed using a modified version of PlotProtein^59^. Candidate residues for generating point mutations to block RNA binding were selected by inspecting the AlphaFold2 models with respect to surface properties. Lysine and arginine residues (K246E, K251E, R539E and K616E), which contribute to positively-charged regions, were identified on the SPRNK domain surface that also corresponded to a region of high conservation across vertebrates.

### Expression and purification of recombinant proteins

Plasmids expressing wild-type and RNA-binding mutant SPRNK domain were transformed into the T7-express BL21 *E.coli*. Cells were cultured in 2x TY broth, and the expression of recombinant proteins was induced by 1 mM IPTG and cultured overnight at 20°C. Cell pellets were lysed using a Continuous High Pressure Cell Disruptor at 4°C and pressure of 25 kPSI. Soluble fractions of protein were purified by ion exchange chromatography on the ACTA Pure system using nickel pre-charged IMAC-HiTrap columns (Cytiva) and the peptides were size separated on a Superdex 200 column. Both, wild type and the RNA-binding mutant were the same size (54kDa) and similar yield.

### Electrophoretic Mobility Shift Assays

To test the binding affinity of the wild-type and RNA-binding SPRNK domain to RNA oligos, and Electrophoretic Mobility Shift Assays (EMSA) were performed. Recombinant protein at increasing concentrations (0–8 μM) were incubated with 0.125 μM fluorophore labeled RNA oligomer (oligoU20-DY681, oligoU40-DY681, oligoA20-DY681, oligoC20-DY681) for 1h on ice in binding buffer (8 mM HEPES pH 7.5, 60 mM KOAc and 16 mM MgOAc) and the reaction products were separated by electrophoresis on 7% acrylamide gel and imaged on the Li-Cor Odyssey instrument. Free and bound RNA were quantified by Image Studio Lite v.5.2.2 software from Li-Cor. Three independent experiments were performed for each condition.

### [5-^3^H]uridine incorporation

Nascent transcription was measured as described previously^16,51^.

### 5-EU incorporation

Cells were treated with 1 mM 5-ethynyl uridine (Base Click) and 1 mM thymidine (Sigma-Aldrich) for the indicated time. Where stated cells were pretreated with 50 ng.ml^-1^ actinomycin D. Click reaction was performed with Click-iT EdU Alexa Fluor 647 Imaging Kit or Click-iT Nascent RNA Capture Kit, according to the manufacturer’s instructions (ThermoFisher).

### Imaging and analysis of 5EU

5EU distribution in nuclei was imaged using a Photometrics Coolsnap HQ2 CCD camera and a Zeiss Axioplan II fluorescence microscope with Plan-neofluar/apochromat objective lenses (Carl Zeiss, Cambridge, UK), a Mercury Halide fluorescent light source (Exfo Excite 120, Excelitas Technologies) and Chroma #83000 triple band pass filter set (Chroma Technology Corp., Rockingham, VT) with the single excitation and emission filters installed in motorised filter wheels (Prior Scientific Instruments, Cambridge, UK). Image capture was performed using Micromanager (Version 1.4).

For the 5EU time-course (Fig. S1G) images were captured using an Andor Dragonfly Spinning Disk Confocal with a 100× objective. 5EU was quantified as follows. Six images containing multiple nuclei were collected for each condition. Nuclei were segmented using the DAPI signal with a home-made script and the mean grey value of 5EU signal was extracted for each nucleus. The mean grey value was normalised to the average mean grey value at 0.5 h (peak value). The difference was calculated between normalised values.

### dSTORM imaging and SuperStructure analysis

#### Immuno-staining for dSTORM imaging

hTERT-RPE1 cells were grown in an 8-well Lab-Tek II Chambered Coverglass #1.5 borosilicate glass (Thermofisher scientific) at 37°C in order to have a final confluency of ∼80%. Cells were fixed with 4% PFA (Sigma-Aldrich) in PBS for 10 minutes, followed by washing in PBS, permeabilisation with 0.2% Triton X-100 (Sigma-Aldrich) for 10 minutes, washed in PBS again, and blocked with 1% BSA (Sigma-Aldrich) for 10 minutes. In the case of SAF-A staining, immuno-fluorescence labelling was done by exposing the cells for 2 h to the primary SAF-A/HNRNPU polyclonal rabbit antibody (A300-690A, Bethyl Laboratories) at 10 μg.ml^-1^ and washed. Cells were incubated for 1 hour with the secondary antibody. The secondary antibody was AffiniPure F(ab’)2 donkey anti-rabbit or donkey anti-mouse IgG (H+L) (711-006-152 and 715-007-003, Jackson ImmunoResearch Europe Ltd) conjugated to the organic fluorophore CF647 (MX647S100, Sigma-Aldrich) at a stoichiometric ratio of about 0.8 fluorophores per antibody molecule.

For RNA staining, 5EU was fluorescently labelled with Cy5 picolyl azide (1 µM) through Click labelling^16^ for 30 min and then washed. When performing two-colour dSTORM where both SAF-A and RNA were labelled, immunofluorescence for SAF-A was performed first and then click labelling for RNA. In this case, the secondary antibody for labelling SAF-A was conjugated to the organic fluorophore CF568 (MX568S100, Sigma-Aldrich). Oxygen scavenger imaging buffer for dSTORM was prepared fresh on the day of use as described^60^. The buffer was prepared as (i) 5.3 ml 200 mM Tris, 50 mM NaCl was mixed with (ii) 2 ml 40% glucose solution, (iii) 200 µl GLOX, (iv) 1.32 ml 1M 2ME (Sigma-Aldrich), and (v) 100ul 50 µg.ml^-1^ DAPI solution (Sigma-Aldrich). The GLOX solution was prepared by mixing 160 µl 200 mM Tris/50 mM NaCl with 40 µl catalase from bovine liver (Sigma-Aldrich) and 18 mg glucose oxidase (Sigma-Aldrich). The final volume was 8.9 ml, which was sufficient to fill the chambers of the 8-well dish; a coverglass was seated on top of the dish to minimise exposure to air.

#### dSTORM Acquisition

STORM acquisitions were performed at room temperature using a Nikon N-STORM system with Eclipse Ti-E inverted microscope with laser TIRF illuminator (Nikon UK Ltd, Kingston Upon Thames, UK). The microscope was equipped with a CFI SR HP Apo TIRF 100× objective lens (N.A. 1.49) with a 1.5× additional optical zoom. A cylindrical astigmatic lens was used to obtain elliptical shapes for emitters that reflect their z-position^61^. Laser light was provided via a Nikon LU-NV laser bed with 405, 488, 561 and 640 nm laser lines. CF647/Cy5 and CF568 fluorophores were stochastically excited using the 640 nm and 561 nm laser lines respectively, with an additional 405 nm weak pulse. A far-red and a multi-colour filter cube were used for single-and two-colour experiments, respectively. Images were acquired with an Andor iXon 897 EMCCD camera (Andor technologies, Belfast UK). The Z position was stabilised during the entire acquisition by the integrated perfect focus system (PFS). For each nucleus a stack of 20,000 frames of 10 or 20 ms exposure time were acquired using the Nikon NIS-Element software. For two-colour dSTORM the far-red (CF647) signal was collected first, and then the red (CF568) signal. Acquired images were 256×256 pixels with equal pixel being 106 nm. For every condition at least 10-15 nuclei (i.e., 10-15 independent datasets) were imaged.

#### Raw images and post-processing analysis

The raw stack of frames was initially segmented based on a DAPI marker to mask out extra-nuclear signal. Frames were analysed using the Thunderstorm plugin^62^ in FIJI. Firstly, frames were filtered by using the Wavelet functions to separate signal from noise. The B-Spline order was set to 3 and the B-Spline scale to 2 for localisations of around 5 pixels. In order to localise the emitter centroids, filtered images were thresholded (threshold value was set to 1.2 times the standard deviation of the 1st Wavelet function) and calculated the local maximum relative to the 8 nearest neighbours. Finally, the emitters signal distribution was fitted to elliptical gaussians (ellipses are necessary for z-position reconstruction) using the weighted least square method and by setting 3 pixels as fitting radius and 2 pixels as sigma. Localised data was then post-processed using the same plugin. (i) XY drift was corrected using a pair correlation analysis, (ii) filtered data with a position uncertainty < 40 nm, (iii) restricted the z-position to the interval [-100:100] nm and projected the data in a 2-dimensional plane, as the z-axis precision is around 100 nm. Reconstructed images shown in the main text were created by using the average shifted histograms method of the Thunderstorm plugin with a 10x magnification (10.6 nm/pixel).

#### SuperStructure analysis for cluster connectivity

dSTORM data was analysed with the SuperStructure algorithm to extract the connectivity between protein clusters^35^. For every nucleus, 5 to 8 local 1.5 µm circular ROIs with were defined and SuperStructure was run within such regions. SuperStructure curves were generated by spanning its spatial parameter *R* ∈ [0,200] nm with a rate *dR* = 10 nm. The super-cluster exponential regime was then visually identified in the averaged curve (as a linear decay in log scale). For both RNA and SAF-A such a regime is *R* ∈ [20,70] nm. SuperStructure regimes were fitted with the exponential function *f*(*R*) = *k* ⋅ *e*^−*R*/*λ*^ and the decay length *λ* was extracted. For every SAF-A simulation, we also calculated the normalised *λ*^∗^ = *λ* ⋅ :*ρ_cl_* to remove the cluster density *ρ_cl_* contribution from the decay length. In the main text, the decay rate *τ* = 1/*λ* is shown for better visual representation, as higher *τ* reflects higher connectivity. The cluster and localisation density were calculated for the same ROIs defined above. Note that in the case of RNA where no evident cluster peak is observed in the Ripley analysis, the localisation density was used instead of the cluster size as observable. Note also that due to spatial inhomogeneity in the RNA signal the SuperStructure analysis was run for the entire nucleus instead of user-defined ROIs.

#### Ripley’s analysis for clustering evaluation

Ripley’s analysis is typically used to verify if a distribution of points is clustered within a certain length-scale, i.e. if it deviates form a random distribution of points. In order to do that, one needs to calculate the average local density of points for different length-scales and properly normalise it. This is the so-called Ripley’s K function:

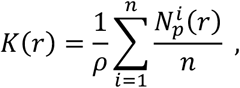

where *N^i^* (*r*) is the number of points found within a distance from a given point *i*, *n* is the total number of points in the system, and *ρ* is the average density of points in the system. The summation is performed for all the points in the system. In the case of a random distribution of points, the density of points is constant and therefore *N^i^* (*r*) = *ρ* ⋅ *πr*^2^. As a result, *K*_rand_(*r*) = *π* ⋅ *r*^2^. If *K*(*r*) > *K*_rand_(*r*), points show a clustering behaviour at that length-scale, otherwise if *K*(*r*) < *K*_rand_(*r*), points are over-dispersed.

The best way to visualise is through Ripley’s H function:

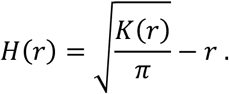

By using this function, *H*_rand_(*r*) = 0 and therefore a clustering phenotype at a given length-scale is represented by a peak of the *H*(*r*) function.

This approach has been used to check the clustering properties of RNA and SAF-A. We performed this analysis on 5 to 8 local square ROIs of 3 *μm* size for every condition. The radius *r* spanned in the range of 0 to 400 nm.

#### Combine DBSCAN and Superstructure for cluster analysis

Cluster analysis was performed by combining SuperStructure with DBSCAN. By visually investigating SuperStructure curves, we identified the value of the spatial parameter *R* to fix for a cluster analysis with DBSCAN. The cluster analysis was performed for that *R* and by fixing the second DBSCAN parameter, i.e. minimum number of localisations to define a cluster, *N_min_* = 0. Finally, the investigation of the distribution of cluster sizes allowed us to define a cut-off to remove clusters below a certain size. In the case of SAF-A, we identified *R* = 20 *nm* and minimum cluster size 30 *localisations*. No cluster analysis was performed for RNA as no evident peak of clustering was observed in the Ripley’s analysis. In that case, the density of localisations was measured.

#### Co-localisation analysis of SAF-A with RNA signal

Co-localisation analysis was performed in sequential steps. Firstly, DBSCAN was run to perform the cluster analysis and extract the centre of protein clusters. Then, a map of the density of RNA in the image was calculated by coarse graining the localisations in pixels of size 10 nm. By doing so a 2D matrix was obtained where each 10 nm bin has the sum of the RNA localisations from STORM raw data. We do this procedure because RNA displays a continuous structure that cannot be easily and unambiguously broken up into clusters. To correlate the SAFA clusters with RNA we employ a method similar to the calculation of radial density function (RDF). While RDF typically computes the presence of a particle at a given distance *r* from another particle, here we compute the cumulative intensity (which we assume proportional to the probability) to find RNA at a given distance from the centre of a SAFA cluster. A 2D map with dimensions 1 μm × 1 μm was compiled, divided in 10 nm bins of the probability of finding RNA at a certain distance in the *x* and *y* directions from SAFA clusters (without assuming the system to be isotropic). This procedure was repeated for all the clusters in each nucleus and across *n* = 10 nuclei. Maps were then averaged to obtain the figures in the main and supplementary figures.

### FRAP and FCS experiments

#### Cell preparation for FRAP and FCS

hTERT-RPE1 cells carrying an inducible copy of GFP multimers and FlpIn T-Rex 293 cells carrying an inducible copy of 3×FLAG–SNAP–SAF-A were grown in 8-well Lab-Tek II Chambered Coverglass #1.5 borosilicate glass (Thermofisher scientific). To label SNAP-SAF-A protein, cells were incubated with SNAP TMR-STAR (NEB) at 1.2 μM for FRAP and 0.06 μM (NEB) for FCS for 30 min and then washed cells in PBS. For acquisition, cell media was replaced with Leibovitz media (ThermoFisher).

#### FRAP acquisitions and analyses

FRAP images were acquired using a Leica SP5 confocal (Leica Microsystems UK Ltd, Milton Keynes, UK) using LAS-AF software with appropriate FRAP wizard. The microscope was equipped with 405 nm diode, Argon and, 561 and 648 nm laser lines, three Photomultiplier tubes and one HyD GaSP detector. Images were scanned using a 40× air objective (N.A. 0.85). Confocal resolution and frequency were respectively set to 128×128bit and 1000Hz in order to obtain a spatial resolution of 179 nm and a time resolution of 140 ms in the Fly-mode of the wizard. Signal of TMR-STAR was excited by using the 561 nm laser at 8% power. FRAP bleaching was performed within a circular region of radius (R) = 2 μm by using all available lasers at 100% power. We acquired 10 pre-bleaching frames (1.4 s), 30 bleaching frames (4.2 s) and 400 post-bleaching frames (56x). For every condition, 20 to 30 FRAP events were acquired on independent nuclei.

FRAP analysis was mainly performed using home-written scripts. Firstly, cell movement was corrected by using the TurboReg plugin in FIJI^63^, then the mean intensity of the bleaching region *I^B^*(*t*) was extracted, as well as that of the entire nucleus *I^C^*(*t*). A first normalisation was performed in order to correct the overall bleaching of the signal in time, as well as to normalise the signal in the range [0,1], where 1 corresponds to the normalised pre-bleaching signal:

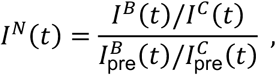

where *I^B^*_pre_(*t*) and *I^C^*_pre_(*t*) are respectively the average pre-bleach signal in the bleached region and in the entire nucleus.

A second normalisation was performed in order to shift the minimum intensity of the signal to 0, while maintaining the signal in the range [0,1]. This is necessary for visually comparing FRAP curves for different conditions, as they typically have different minima in the intensity (bleaching is not 100% efficient). Note, that this second normalisation does not influence the fitting of the curves. The new normalised intensity is therefore

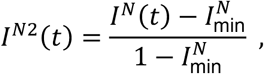

with being *I^N^* the minimum intensity of the FRAP curve. The recovery of FRAP curves (*I^N^*^2^(*t*)) was fitted with an exponential function by assuming the existence of two mobile populations (fast and slow population)

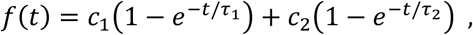

where *τ*_1_ and *τ*_2_ are respectively the recovery time of the fast and slow population. These two times can be though as inversely proportional to the diffusion time of that population: *D*∼*R*^2^/*τ*. The fraction of the mobile population is defined as *MF* = *c*_1_ + *c*_2_. As a consequence, one defines the fraction of signal that is not recovered as the immobile fraction *IF* = 1 − *MF*. Note that *IF* can be defined also independently from the fitting function, by measuring the fraction of signal that is not being recovered.

#### FCS acquisitions and analyses

For SNAP–SAF-A, FCS data was acquired using Leica SP5 Confocal with appropriate wizard using a PicoHarp 300 TCSPC module and APD detectors (Picoquant). Images were scanned using a 40X water objective. Confocal resolution and frequency were respectively set to 256×256bit and 600Hz. Signal of TMR-STAR was excited by using a 561 laser. FCS acquisition was performed in a very small volume and we measured the intensity signal for 100 s. For every condition, ∼50 FCS measurements were acquired on independent nuclei.

For GFP multimers, data was acquired with a Zeiss LSM880 equipped with the FCS module (Carl Zeiss, Jena, Germany). GFP was excited with a 488-nm Argon laser line and scanned using a 63× (N.A. 1.2) water immersion objective (Zeiss). To calibrate the size of the point spread function, a 100 nM solution of Rhodamine 6G was prepared in PBS buffer and the beam waist of the resulting data was fitted.

The acquired data was processed using Symphotime 2.6 or Zen 2.3 SP1. The signal was restricted within an appropriate time-window where no bleaching was observed (10s in case of SNAP–SAF-A). We calculated the autocorrelation of the signal as function of the time lag

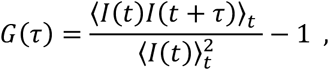

where *I*(*t*) is the signal intensity at time *t* and (⋅⟩*_t_* is the sliding average over all values of *t*. We then fitted the autocorrelation curve with a single-population free diffusion model^64^:

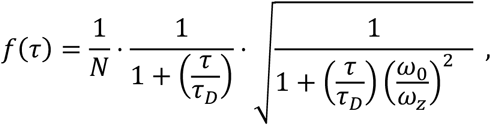

where the model contains three fitting parameters: the average number of particles inside the confocal volume *N*, the structure parameter *κ* = *ω*_0_/*ω_z_* that describes the shape of the effective detection volume and the proteins’ diffusion time *τ_D_*. The values of *ω*_0_ and *ω_z_* are determined from a calibration with free-diffusing dye with known diffusion coefficient. Finally, the diffusion coefficient of proteins is calculated as *D* = ω_0_^2^/4τ_*D*_.

### SAF-A oligomerization assay and detection

SAF-A oligomer formation was analysed as described previously^16^.

### BioID2 assay

FlpIn T-REx 293 cells carrying an inducible copy of myc–BioID2–SAF-A or myc–BioID2 as negative control were incubated in the presence of 50 μM biotin for 24 hours. After collecting cells with trypsin treatment and PBS wash 3 times to remove free biotin, the cell pellet (corresponding to 2 × 10^7^ cells) was lysed in 500 μL of RIPA Lysis buffer (Santa Cruz Biotechnology, sc-24948) and sonicated to reduce sample viscosity from lysed DNA. Biotinylated proteins were purified using streptavidin magnetic beads and subjected to proteomics analysis by the IGC mass spectrometry facility. Samples were Strep Precipitated and processed on a Kingsfisher Duo robot. IPs, washes and on-bead digests were performed using a Thermo Kingfisher Duo, all steps are at 5°C unless otherwise stated. Beads were transferred into 500 μl of cleared lysate and incubated for 2 h with mixing. Beads were transferred for two washes in RIPA buffer and three washes in non-detergent lysis buffer. On-bead digest was performed by transferring the washed beads into 100 μl 2M urea, 100 mM Tris-HCl pH 7.5, 1 mM DTT containing 0.3 μg trypsin (Promega) per sample, beads were incubated at 27°C for 30 min with mixing to achieve limited proteolysis. The beads were then removed and tryptic digest of the released peptides was allowed to continue for 9 h at 37°C. Reduced cysteine residues were alkylated by adding iodoacetamide solution to a final concentration of 50 mM and incubated 30 min at room temperature, in the dark. Trypsin activity was inhibited by acidification of samples to a concentration of 1% TFA. Samples were desalted on a C18 Stage tip and eluates were analysed by HPLC coupled to a Q-Exactive mass spectrometer as described in^65^.

Peptides and proteins were identified and quantified with the MaxQuant software package (1.5.3.8), and label-free quantification was performed by MaxLFQ^66^. The search included variable modifications for oxidation of methionine, protein N-terminal acetylation, and carbamidomethylation as fixed modification. The FDR, determined by searching a reverse database, was set at 0.01 for both peptides and proteins.

### Electron microscopy

293T cells expressing FLAG–SAF-A were suspended in ATPase buffer in the presence or absence of RNaseA/T1 for 10 min at 37°C. Samples were treated with BM(PEG)2 protein cross-linker for 5 min at RT [as described previously^16^] and quenched using L-cysteine. Following sonication to break genomic DNA, samples were clarified by centrifugation. Supernatant was incubated with FLAG-agarose beads for 2 h at 4°C and SAF-A was eluted with 3 × FLAG peptide. For TEM, samples were absorbed on coated grids and negatively stained with 2% uranyl acetate and visualised using a Philips/FEI CM120 Biotwin microscope.

### Molecular dynamics simulations of chromatin and RNA–SAF-A microgels

The structure formed by chromatin and RNA–SAF-A mesh of microgels is investigated by performing molecular dynamics simulations. These evolve the equation of motion of explicitly resolved chromatin (as a bead-and-spring polymer) and SAF-A (as patchy particles), within an implicit heat bath that mimics the viscous nuclear environment. In our polymer model, we consider a chromatin stretch with multiple active euchromatic and inert heterochromatic regions. The beads in the active segments act as a source for an RNA field that triggers SAF-A oligomerisation. Such oligomerisation creates microgels which can further be interconnected, according to the system parameters. The system undergoes a sol-gel transition: in the transition region and the gel phase we observe the decompaction of the active chromatin regions, as shown in experiments.

#### Polymer model for the chromatin fibre

The chromatin is modelled as a bead-spring polymer chain where every bead has a nominal diameter *σ* which corresponds to 1 kbp and to about 20 nm in size (with the exact value determined by fit to SuperStructure microscopy data, see below), in the range used in our previous coarse-grained models for chromatin^67^.

In our present model, the chromatin fibre consists of a concatenation of ten active (euchromatin) domains, and ten inactive (heterochromatin) domains, regularly spaced, and with size 200 beads (200 kbp, for euchromatin) and 300 beads (for heterochromatin). In euchromatin segments, there are equally spaced highly sticky beads, which represent regulatory units (promoters or enhancers): their spacing was set to 20 beads, starting from the 5-th bead in each domain.

The connectivity of the entire chain is provided by a harmonic potential between consecutive beads (i.e., a spring with large stiffness):

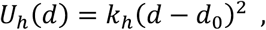

where *d* is the distance between two consecutive beads, *k*_ℎ_ = 200*k_B_T*/*σ*^2^ and *d*_0_ = 1.1*σ*. Note that *k_B_T* is the energy unit, where *k_B_* is the Boltzmann constant and *T* the temperature.

The chromatin chain is also subject to a bending potential to introduce a persistence length (or bending stiffness) of the fibre:

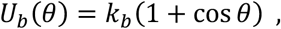

where *θ* is the angle between two consecutive bonds and *k_b_* = 3*k_B_T*. Such parameters allow to mimic a persistence length *l_p_* ≃ 3*σ*.

An additional potential between chromatin beads is set to (implicitly) mimic protein bridges which drive folding and compaction of chromatin, as well as to introduce steric interaction between beads. This is a cut-and-shifted Lennard-Jones potential:

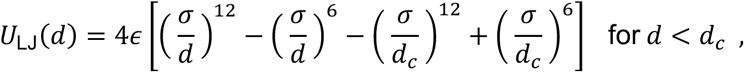

and 0 elsewhere. We chose (i) *ε* = 10 *k_B_T* and *d_c_* = 2.5*σ* between beads corresponding to regulatory elements, to set up strong attraction between promoters and enhancers, and (ii) *ε* = 0.7 *k_B_T* and *d_c_* = 2.5*σ* to set up a weak self-attraction potential between active chromatin beads which are not promoters. For all other types of chromatin beads, we set the parameters *ε* = *k_B_T* and *d_c_* = 2^1/6σ^, which model purely steric interactions. Finally, we add one patch on each bead along chromatin. The patches on the active chromatin and regulatory element beads are considered to be newly synthesised RNA and can bind to those on SAF-A (see below), whereas patches on heterochromatin are inert.

#### The equation for the RNA field

The beads of the active gene locus can produce newly transcribed RNA at a rate *k*_prod_ and with a probability of 10%. Newly transcribed RNA then diffuses with a (small) coefficient *D*, and is degraded at rate *k*_deg_. Instead of explicitly simulating each RNA molecule in the system (which would be too expensive computationally), we solve the diffusion equation associated with the dynamics of the RNA density field. Given an RNA-producing source at location ***r***_0_ (i.e., a bead of the gene locus), the density of RNA signal *ρ*(***r***, *t*) evolves in space and time as the following reaction-diffusion equation

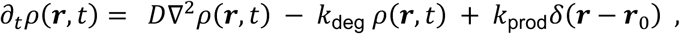

where the terms on the right side of the equation are respectively the diffusion, the degradation and the (localised) production terms. The function *δ*(***r*** − ***r***_0_) is the Dirac delta function.

Since we perform simulations at equilibrium (or near equilibrium condition), we can consider the steady state form of the equation, i.e., *∂_t_ρ*(***r***, *t*) = 0. The resulting equation is also known as screened Poisson, or Yukawa, equation, and can be solved with the method of Green’s functions. In other words, one needs to solve

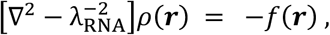

where *f*(***r***) = *k*_on_*δ*(***r*** − ***r***_0_)/*D* is the source term and λ_RNΑ_ = :*D*/*k*_deg_ is the RNA decay length, which measures the typical distance over which RNA diffuses before being degraded in our case. This equation is solved by first finding the associated Green’s function *G*(***r***) that is the solution of the equation

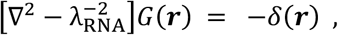

which, in Fourier space, reads

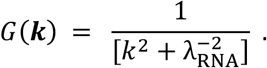

In 3D, the inverse transform gives

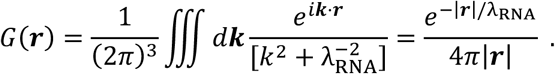

Hence, the RNA field in real space is found as

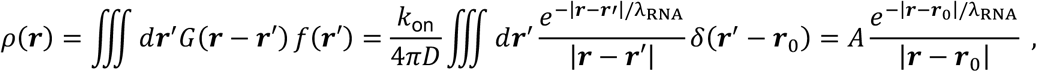

where *A* = *k*_prod_/4*πD* can be viewed as the gene activity. This equation shows that the contribution to the concentration of RNA at position ***r*** is given by a factor proportional to how much RNA is produced by the active locus (*A*), multiplied by a screening Yukawa function that depends on how much RNA is degraded before reaching ***r*** by diffusion (*λ*). In virtue of the superposition principle, if we have *n* independent sources of RNA, the contribution to location ***r*** is the linear sum of contributions of the *n* sources. Therefore, thanks to this equation we can calculate the instantaneous (steady-state) local concentration of newly synthesised RNA given the position of the *n* sources of RNA. The number of RNA molecules in the locus may be found by integrating the RNA density over a volume enclosing the gene locus.

#### Polymer model for SAF-A proteins

SAF-A proteins are simulated as diffusing beads in the space. Steric interactions (with either chromatin beads or other proteins) are considered through a Lennard-Jones potential similar to that used between chromatin beads, with *ε* = *k_B_T* and *d_c_* = 2^1/6^ *σ*. In order to implement the RNA-binding properties of SAF-A, each protein bead is enriched with two additional patches. First, we introduced an attractive interaction between SAF-A patches and those on active chromatin and regulatory element beads (representing newly synthesised RNA) via a Morse potential:

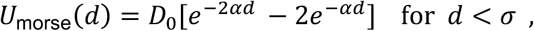

where *D*_0_ = 30*k_B_T* for active chromatin beads and *D*_0_ = 50*k_B_T* for regulatory element beads, while *α* = 16*σ*^−1^. The idea underlying this potential choice is that SAF-A coats the active chromatin region, binding to its beads via RNA. Some form of scaffolding to the chromatin polymer is required in order to ensure that the self-assembled SAF-A/RNA microgel remains in the vicinity of transcriptionally active chromatin, without being entropically pushed away by the polymer.

Second, to implement the formation of the RNA–SAF-A mesh, we considered a two-state model for SAF-A beads. Thus, any of the SAF-A proteins can be either in an on (RNA-bound) or off (not RNA-bound) state. The conversion rate *k*_on_ from the off to the on state is set to be proportional to the local (calculated) concentration of RNA *ρ*(***r***), whereas the inverse, from the on to the off state, is set to be a constant turnover rate *k*_off_. Thus, at any given timestep of the simulation, there is a subset of SAFA proteins in the on state, and another in the off state. When the protein is in the *on* state, the two patches can bind to the patches of other on SAF-A proteins via the same Morse potential introduced above. The values of the parameters in the Morse potential between SAF-A patches are the same as those between SAF-A and chromatin patches: they lead to the creation of frequent oligomers with about 1 branch in 5 to10 monomers. Despite of the large energy strength set for this potential, the self-assembled structure is still dynamic due to the non-equilibrium switching to the off state at rate *k*_off_.

#### Molecular dynamics simulations

The dynamics of each chromatin and protein bead *i* with mass *m* evolves according to a Langevin equation at constant temperature *T*:

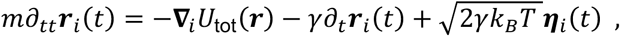

where *U*_tot_ is the total potential energy resulting from the addition of the above interaction terms, **∇***_i_* is the gradient taken respect to the coordinates of the *i*th bead, *γ* is the friction coefficient and ***η****_i_* is the delta-correlated stochastic white noise acting on every bead. Note that every SAF-A bead with its 2 patches is defined as a rigid body with mass *m*_prot_ = 3*m*, while every chromatin bead is defined as a rigid body with mass *m*_chr_ = 2*m*. We set the friction coefficient equal to *γ* = *m*_chr_/2*τ*_LJ_ for each chromatin bead + patch and to *γ* = *m*_prot_/3*τ*_LJ_ for each protein bead + 2 patches, where *τ*_LJ_ = *σ*:*m*/*k_B_T* is the Lennard-Jones time. This choice ensures that both polymer and protein beads in isolation (i.e. not bound or joined into a chain) would have the same diffusion coefficient (even though they are formed by 2 and 3 particles, respectively). With this choice, the typical simulation time is *τ*_LJ_ = 3*πη_s_σ*^3^/*k_B_T* ≃ 1 ms for *σ* = 20 nm and with the viscosity of the nucleus set to be *η_s_* = 200 cP ^68^. The equation of motion is integrated with timestep Δ*t* = 0.002*τ*_LJ_ using the LAMMPS simulation package ^69^.

#### Simulation parameters

In our standard simulations we consider a chain of 5000 chromatin beads, organised as ten concatenated pairs of a 200-bead active and a 300 inactive domain. By assuming that each chromatin bead is around 1 kbp, the active domains in our simulations are each 200 kbp, whereas the whole fibre simulated is 5 Mbp long. We consider 4000 SAF-A proteins diffusing in proximity of the chain with an overall protein volume fraction of 0.033, or a protein density of 0.0625*σ*^−3^. F or what regards RNA, we vary λ_RNΑ_ in the range of 1*σ* to 30*σ*, and *A* between 0.01 and 1.8*σ*^−2^. In terms of the dynamics for the RNA field (and given a value of *σ* = 20 nm) these can be mapped, for instance, to a range of turnover rates for RNA *k*_deg_ between 0.0027 and 2.5 s^−1^, an RNA diffusion coefficient *D* ≃ 10^−3^ μm^2^/s and a range rate of RNA production *k*_prod_ ≃ 0.31 to 57 s^−1^. For the SAF-A modelling, the SAF-A de-oligomerisation rate is set to *k*_off_ = 1/*t*_off_, with *t*_off_ = 600 *τ*_LJ_ = 0.6 s, whereas the oligomerisation rate is *k*_on_ = *k*_0_min {1, *ρ*}, with *k*_0_ = 0.01 *τ*^−1^, and we recall *ρ* denotes the RNA density field.

#### Simulations with different parameters and comparison with experiments

To generate simulations to be compared with experiments, we varied *A* in the range of 0.1*σ*^−2^ to 1.8*σ*^−2^ and λ_RNΑ_ in the range of 1*σ* to 5*σ*. We also performed selected simulations with *A* in the range [0.01, 1] *σ*^−2^ and λ_RNΑ_ = 30*σ*. For each simulation, we computed simulated SuperStructure curves, and fitted them by allowing a variation of the size of a simulation bead between 20-25 nm, and an arbitrary multiplicative factor rescaling of the number of cluster per localisation (note this does not affect the slope in the log/linear curve we measure). For each fit, we measured the value of *χ*^2^, and found that it is minimised (corresponding to the best fits) in the transition regime. We also computed the FISH expansion ratio, measured as the average distance between beads in active euchromatin regions at separation along the chain between 120-150 beads, normalised with the values corresponding to the same separations recorded for (*A*, λ_RNΑ_) = (0.1,1). These normalised distances tend to increase with both *A* and λ_RNΑ_, and hence give a measure of the decompaction induced by transcriptional activity.

#### FCS simulations

To predict 3D diffusivity and mobility of GFP multimers observed experimentally by FCS, we simulated 3D diffusion and mobility of beads of variable size (diameter) *σ*_GFP_, with 0.4*σ* ≤ *σ*_GFP_ ≤ 2*σ*. These GFP spheres diffused through a static chromatin–SAF-A mesh, which, for given values of (*A*, λ_RNΑ_), was made up by the chromatin and active SAF-A beads in the final snapshot of a coarse-grained polymer simulation (see above) with these parameters. We simulated multiple sizes and multiple spheres for each size in parallel in order to increase statistics. While we included steric interactions between the GFP spheres and the mesh, via WCA potentials with *σ* replaced by (*σ* + *σ*_GFP_)⁄2 in the definition of the potential (see above), we did not include steric interactions between GFP spheres, to reflect the experimentally relevant condition in which GFP multimer act as low-density probes of nuclear viscosity. To compare our simulations to FCS experiments, we have considered the case in which the free diffusion of a GFP sphere of size 20 nm is given by ∼20 µm^2^/s, corresponding to a background aqueous viscosity of 1 cP without any polymer or SAF-A gel.

**Fig. S1. Properties of nuclear RNAs.** (**A**) Time course of [5-^3^H]uridine pulse labelling of RNA in RPE1 cells. Actinomycin D was added to cells before a 15 min pulse of [5-^3^H]uridine, followed by sampling over a 2 h period. (**B**) Quantification of newly transcribed RNA in the cytoplasm and nucleus of cells pulse labelled with [5-^3^H]uridine for 15 min in absence (top panel) or presence (bottom panel) of low dose actinomycin D, to inhibit RNA polymerase I. (**C**) (Left) Agarose gel showing the size distribution of total nuclear RNA, stained using ethidium bromide and imaged on a laser scanner, after pulse labelling with [5-^3^H]uridine in the absence of actinomycin D. (Right) Gel shown in left panel transferred to nylon membrane, and analysed using a phosphor imager to detect newly transcribed RNA. (**D**) Fluorescence imaging of cells after pulse labelling with 5EU for 15 min to mark newly transcribed RNA, in the presence of actinomycin D. 5EU was detected using click chemistry with an azide-conjugated fluorescein molecule. (**E**) Fluorescence imaging of cells continuously labelled with 5EU for 24 h. After incubation, samples were fixed and RNA was detected using click chemistry with an azide-conjugated fluorescein moiety and imaged using a wide-field microscope. (**F**) Time course of 5EU labelling of newly synthesised RNA. Samples were collected at 0, 0.5, 1, and 3 h. (**G**) (Left) Fluorescent imaging of cells after pulse labelling with 5EU to mark newly synthesised RNA. After fixation RNA was detected using click chemistry with an azide-conjugated fluorescein moiety and imaged using a Spinning Disk Confocal. (Right) Nuclear RNA quantification. *p* values for Student’s *t*-test: **, p < 0.01; ****, p < 0.0001 (n = 20). (**H**) (Left) two-colour STORM imaging of SAF-A (yellow, CF568 dye) and RNA (purple, CF647 dye), at different times (0, 0.5, and 1 h) after RNA labelling. For images showing RNA (left panels), inset corresponds to enlarged region marked by dashed box. For RNA/SAF-A images (middle panels), dashed box corresponds to enlarged view shown in right panel. Blue arrows correspond to regions of pronounced SAF-A and RNA overlap. (Right) Meta-analysis of RNA signal strength surrounding SAF-A clusters (n = 10 nuclei).

**Fig. S2. SPRNK domain structure and RNA binding characteristics.** (**A**) Cartoon of the SAF-A protein depicting domains and their secondary protein structure (alpha helix, dark blue; beta strand, light blue). Lollipops indicate sites of post-translational modifications. Protein conservation (calculated by ConSurf) is shown from low (turquoise) to high (purple); positions of benign missense variants (from gnomAD) are dark blue lines. Vd/Vp ratios of missense variant depletion per domain compared to depletion over the full-length protein are shown below domains. The SPRNK domain construct (SPRY/PNK) used in binding experiments is shown. (**B**) AlphaFold2 models of SAF-A SPRNK domain showing secondary structure, electrostatic charge (APBS) and protein conservation (CONSURF). (**C**) Graph showing melting temperature of SPRNK domain in the presence of different concentrations of ATPγS. (**D**) Acrylamide gel of an electrophoretic mobility shift assay (EMSA) with fluorescently labelled RNA of either 20 or 40 nucleotides and purified SAF-A SPRNK domain. (**E**) EMSA with fluorescently labelled RNA consisting of either oligo-A, oligo-C or oligo-U with purified SAF-A SPRNK domain. (**F**) EMSA with fluorescently labelled RNA and purified SPRNK domain in the presence or absence of nucleotides. Right, cartoons depicting the potential interactions between SAF-A, RNA and ATP. (**G**) Top, cartoon showing the SAF-A SPRNK domain with putative residues marked for RNA binding. Bottom, putative RNA binding residues superimposed on the secondary structure of SPRNK domain. (**H**) EMSA with fluorescent RNA template with wild-type (WT) or putative SPRNK RNA-binding mutant. (**I**) Quantification of EMSA. P-values are for a T-test, n=3. (**J**) Transmission electron microscopy (TEM) of negatively stained immuno-purified FLAG-tagged full length SAF-A prepared in the presence and absence of RNAseA/T1. Scale bars, 500 nm. (**K**) Boxplot quantifying the length of the major and minor axis of SAF-A particles shown in D (n>50 particles; P values are for a Wilcoxon test; ****, p < 0.0001).

**Fig. S3. Superstructure algorithm, and curves for SAF-A and RNA.** (**A**) Left, cartoon showing how cluster analysis with an increasing radius is built, and a typical shape. Right, cartoon showing how to interpret the resulting SuperStructure decay curve to infer network properties and quantify network connectivity. A low decay rate indicates low connectivity, whilst a high decay rate suggests high connectivity. (**B**) SuperStructure connectivity decay curves for SAF-A and for the connectivity decay curves for newly synthesised RNA matured for 0.5 h. (**C**) SuperStructure connectivity decay curves for newly synthesised RNA matured for 0 h, 0.5 h, and 1 h.

**Fig. S4. Superstructure curves for RNA after SAF-A depletion.** SuperStructure connectivity decay curves for RNA after SAF-A depletion by siRNA. Control, black line; SAF-A depletion, blue line.

**Fig. S5. Microscopy of RNA**–**SAF-A microgels following transcriptional inhibition.** (**A**) Top, time course of transcription inhibition in RPE1 cells. Actinomycin D was added to cells, after 1 h 5EU was added followed by the addition of transcription inhibitor (triptolide or α-amanitin), with sampling at 4 h. Bottom, Imaging 5EU labelled RNA by conjugation to Alexa488 using Click chemistry before and after transcription inhibition. Scale bar is 10 μm. (**B**) Top, time course of transcription inhibition in RPE1 cells. Actinomycin D was added to cells, after 45 min [5-^3^H]uridine was added followed by the addition of α-amanitin, with sampling over a 3 h period. Middle, agarose gel of RNA isolated from cells pulse labelled with [5-^3^H]uridine in the presence of absence of α-amanitin, stained with ethidium bromide. Markers correspond to 28S and 5S RNA. Bottom, newly synthesised radioactive RNA from gels above, blotted onto membrane and analysed on a phosphor imager. (**C**) Top, boxplot showing SAF-A localisation density imaged using STORM for control and triptolide treated cells (n > 60 regions). Bottom, SAF-A cluster density imaged using STORM for control and triptolide treated cells (n > 60 regions). (**D**) SuperStructure connectivity decay curves for SAF-A after transcription inhibition with triptolide. Control, black line; triptolide treatment, red line.

**Fig. S6. Identification of XRN2 and role in determining nuclear RNA turnover.** (**A**) Table showing proteins interacting with SAF-A by BioID. Proteins are organized and coloured in families. These data show that the appearance of XRN2 as a key interacting partner of SAF-A. (B) Cartoon depicting a possible mechanism for the turnover of RNA–SAF-A microgels.

**Fig. S7. Characterisation of nuclear RNA turnover.** (**A**) Top, Diagram explaining how experiment was performed. Bottom, fluorescent imaging of 5EU in newly synthesised RNA labelled by click chemistry with an azide-Alexa488 dye, in control cells or cells depleted for EXOSC3 (exosome) or XRN2. Right, quantification of 5EU in control or XRN2 depleted cells over time (n = 10 nuclei). (**A**) Dot blot showing quantification of newly transcribed nuclear and cytoplasmic RNA labelled with 5EU, across a 6 h period. (**B**) Top, immunofluorescence staining of XRN1 and XRN2 in control and ribonuclease depleted RPE1 cells. Bottom, testing specificity of XRN1 and XRN2 antibodies and siRNAs in RPE1 total cell extracts after treatment with either control, XRN1 or XRN2 siRNAs. (**C**) Top, immunofluorescence staining of EXOSC3 (exosome subunit) in siCTRL and siEXOSC3. Bottom, characterisation of EXOSC3 antibody by Western blotting in control or EXOSC3 depleted cells. (**D**) Western blot for XRN2 degron in HCT116 cells after auxin treatment. (**E**) Detection of 5EU incorporation and turnover in newly synthesised RNA by conjugation to fluorescent Alexa488 in control and auxin treated cells.

**Fig. S8. XRN2 depletion following triptolide treatment increases microgel connectivity.** (**A**) SuperStructure connectivity decay curves for SAF-A after XRN2 depletion. Control, black line; siXRN2, orange line. (**B**) SAF-A STORM in triptolide treated cells with or without XRN2 depletion. Scale bar is 250 nm. (**C**) SuperStructure decay curve of the connectivity between RPE1 cells which were treated with the transcription inhibitor, triptolide, before (siCTRL+Triptolide) and after (siXRN2+Triptolide) XNR2 depletion (n > 90 regions). P values are for a T-test; ns, not significant; *, p < 0.05; **, p < 0.01; ***, p < 0.001; ****, p < 0.0001.

**Fig. S9. Live cell experiments provide an estimate of protein mobility.** (**A**) Top, analysis of HALO-SAF-A by fluorescence correlation spectroscopy. Representative autocorrelation analysis for control cells and for cells treated with the transcription inhibitor, triptolide. Bottom, boxplots comparing the diffusion coefficient for SAF-A in control cells, and cells treated with triptolide. (**B**) Top, representative SAF-A FRAP curves and bottom, SAF-A fast population recovery times for control, triptolide treated or triptolide treated and XRN2 depleted cells (n > 20 regions). (**C**) Cartoon, showing structure and genomic position of different GFP constructs at the ROSA26 locus. (**D**). Fluorescent images of GFP constructs in RPE1 cells before and after induction with doxycycline. P values are for a T-test; ns, not significant; *, p < 0.05; **, p < 0.01; ***, p < 0.001; ****, p < 0.0001.

**Movie S1:** Movie of the trajectory of the coarse-grained molecular dynamics simulations corresponding to the sol phase and the snapshot in Fig. 2C, with *A* = 0.75 × 10^−3^ nm^−2^ and *λ_RNA_* = 40 nm. Left: active SAF-A (green) is shown together with euchromatin (orange); genes in euchromatin are depicted in blue, and heterochromatin is not shown for visual clarity. Middle: active SAF-A only. Right: isosurfaces of the SAF-A density (blue), which are here shown as a proxy for SAF-A clusters. In the simulation there are 5000 chromatin beads and 4000 SAF-A proteins in total (this includes both active and inactive).

**Movie S2:** Movie of the trajectory of the coarse-grained molecular dynamics simulations corresponding to the gel phase and the snapshot in Fig. 2D, with *A* = 3.75 × 10^−3^ nm^−2^ and *λ_RNA_* = 80 nm. Panels and other parameters are as for Movie S1.

**Movie S3:** Movie of the trajectory of the coarse-grained molecular dynamics simulations corresponding to the transition regime and the snapshot in Fig. 2G, with *A* = 2.5 × 10^−3^ nm^−2^ and *λ_RNA_* = 60 nm. Panels and other parameters are as for Movie S1, except for the number of total SAF-A proteins, which here is 2000.

**Movie S4:** Trajectory of an 8 nm sphere in the gel phase, corresponding to high transcriptional activity. The left panel shows the trajectory of the sphere together with the SAF-A/chromatin gel, with chromatin (thick polymer) and active SAF-A (beads), whereas the right panel shows the trajectory only for visual clarity. It can be seen that smaller particles are transiently trapped inside the gel.

**Movie S5:** Trajectory of a 40 nm sphere in the gel phase, corresponding to high transcriptional activity. The left and right panels are as in Movie S4. It can be seen that larger particles are mostly excluded from the gel.

