## Supplementary material for "A transcription-driven microgel network poised near the sol-gel transition": Supp figures

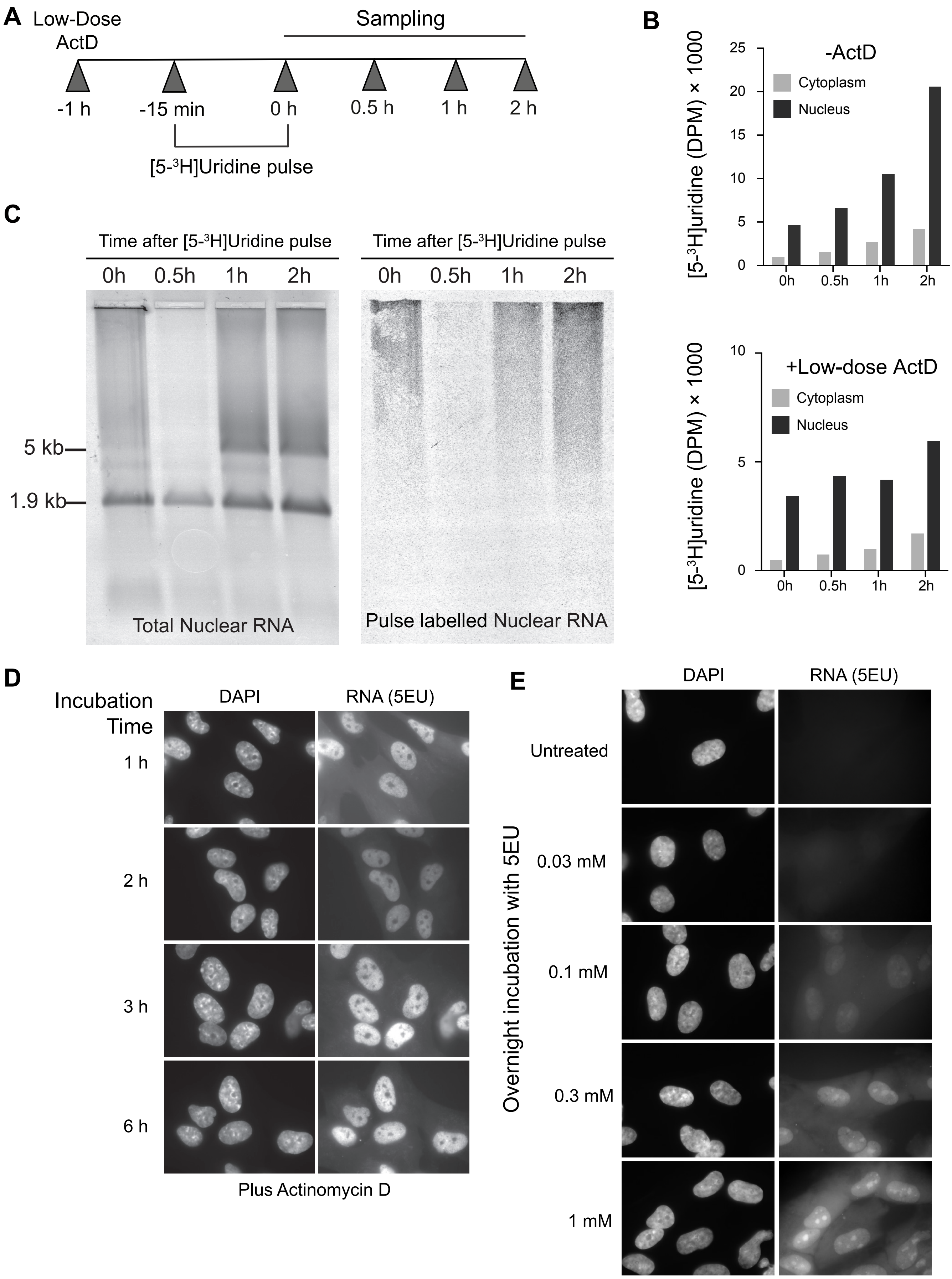


Marenda et al. Supplementary Figure 1

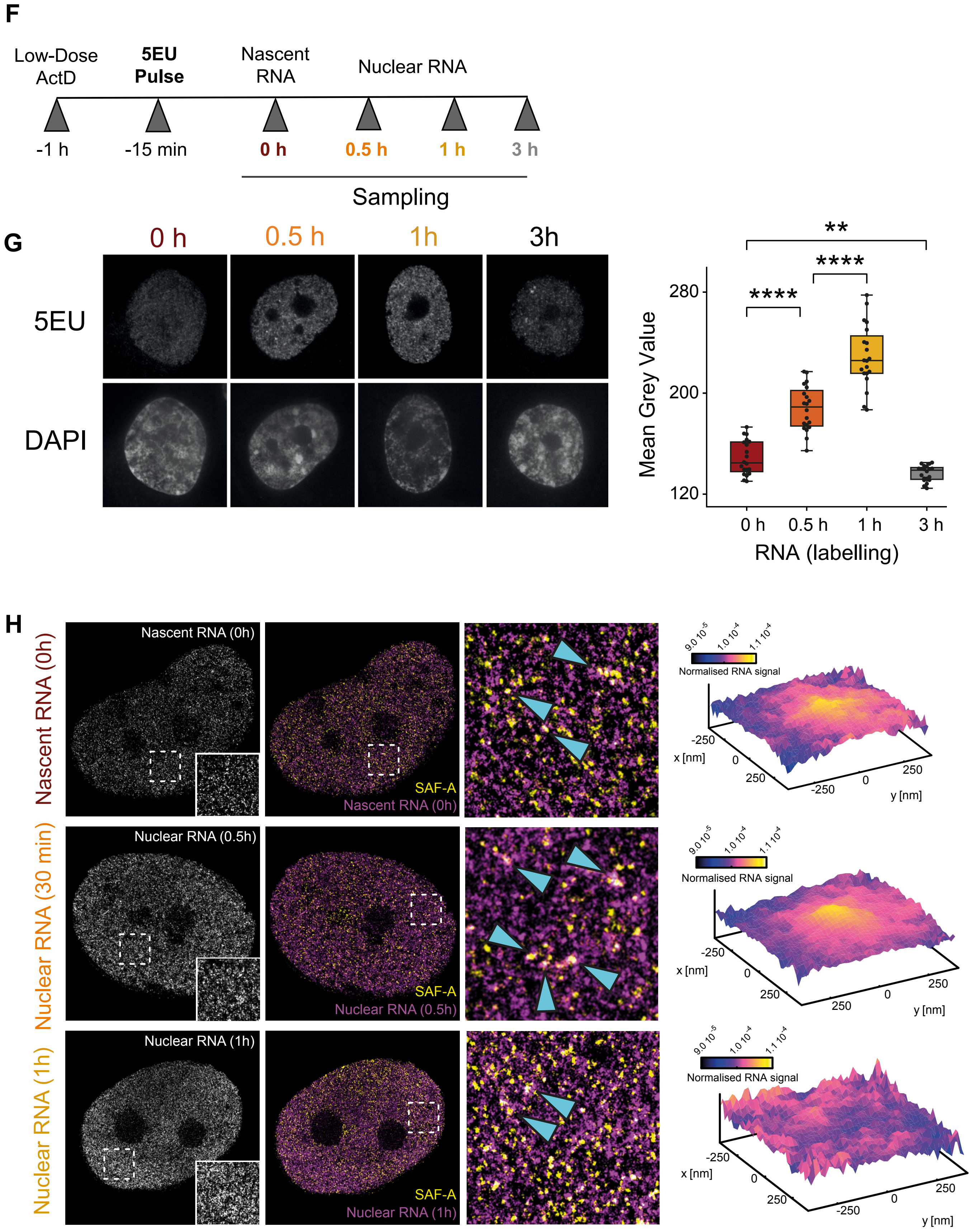


Marenda et al. Supplementary Figure 1

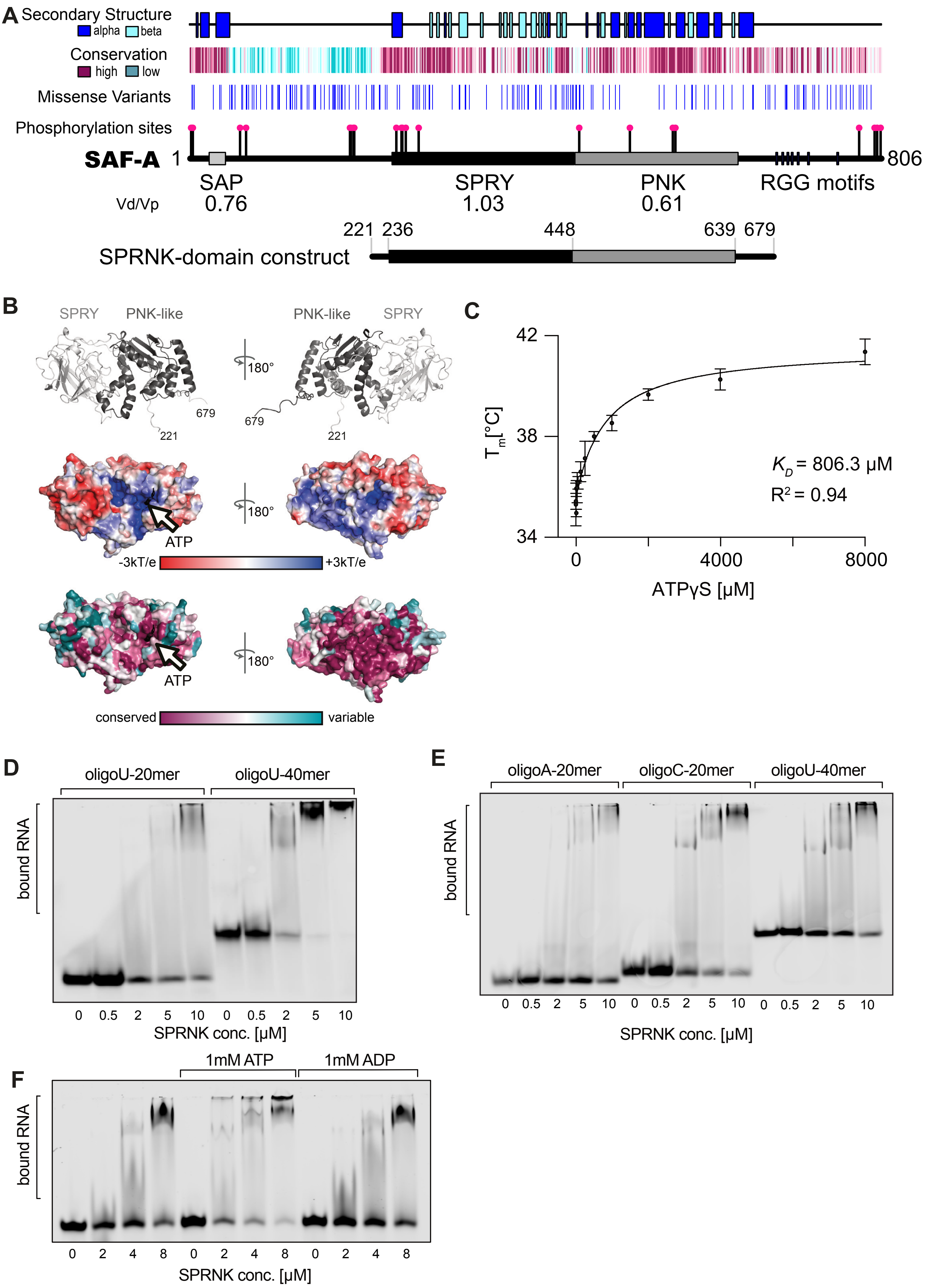


Marenda et al. Supplementary Figure 2

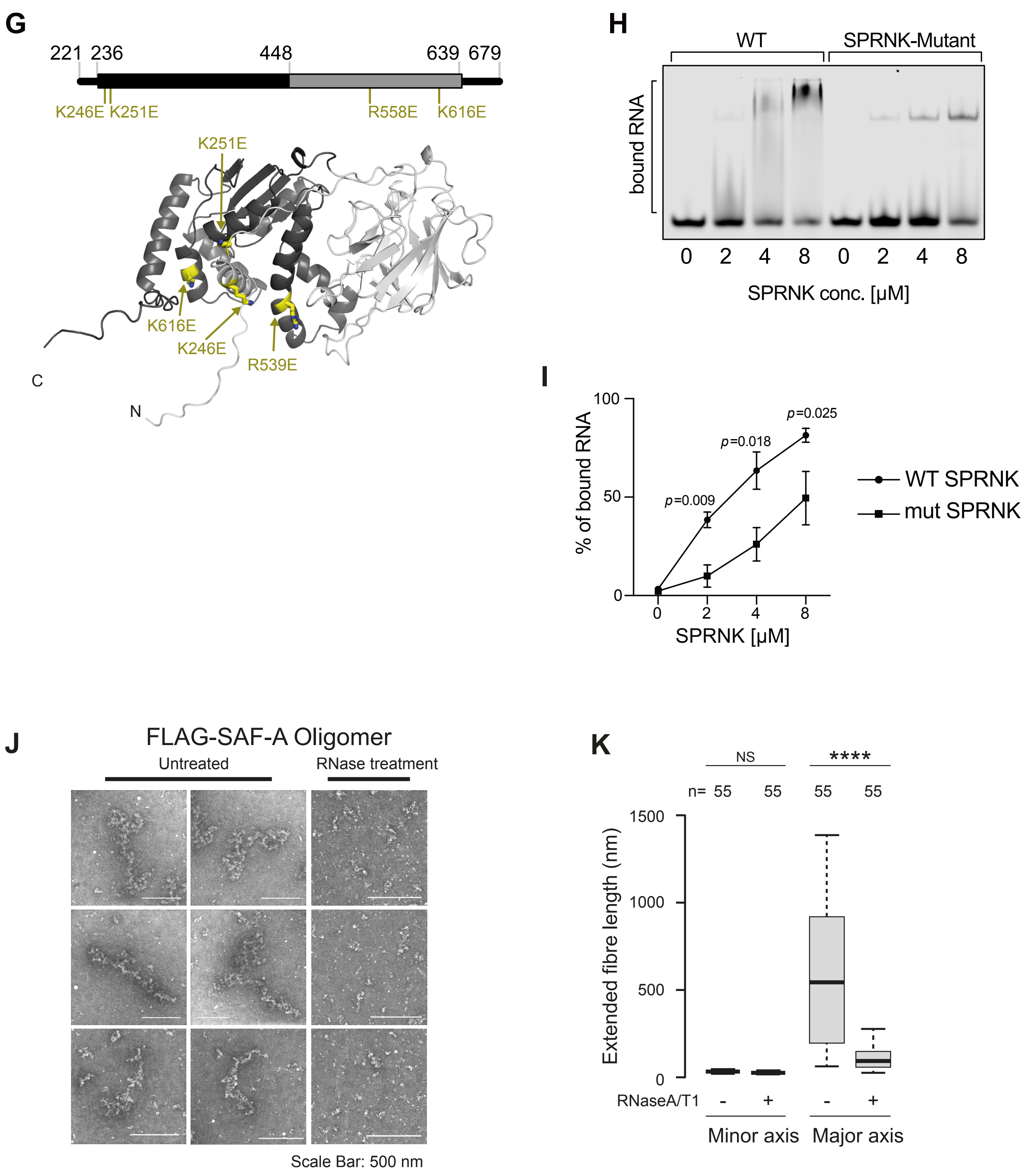


Marenda et al. Supplementary Figure 2

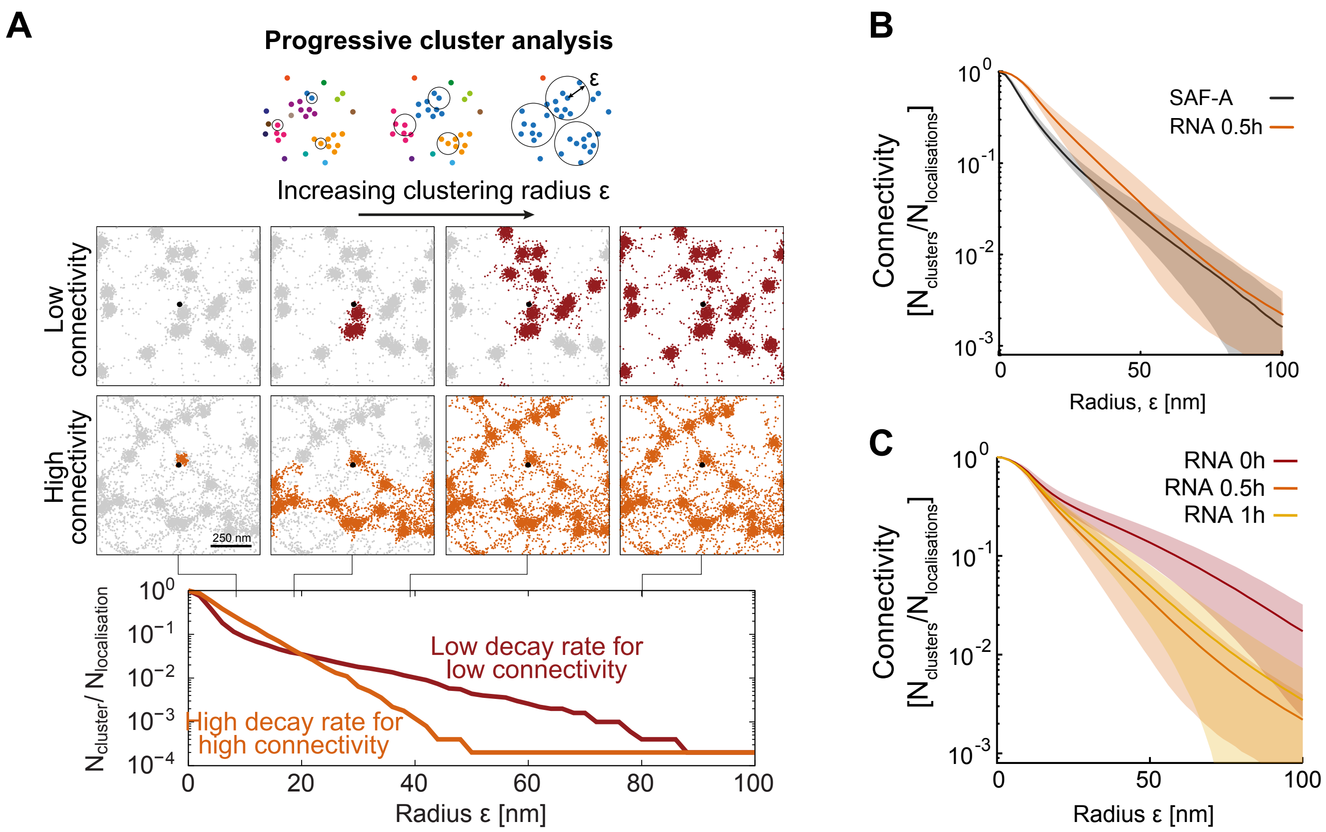


Marenda et al. Supplementary Figure 3

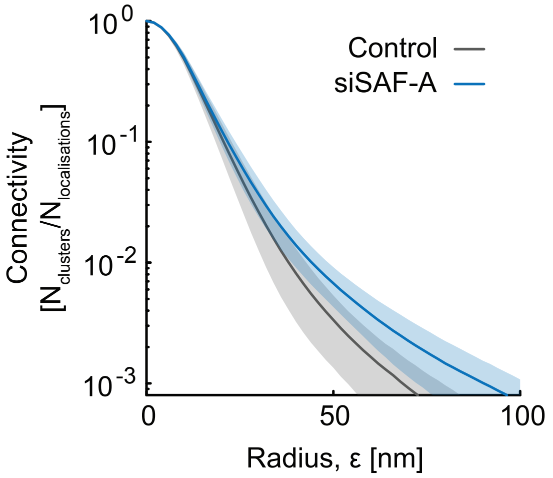


Marenda et al. Supplementary Figure 4

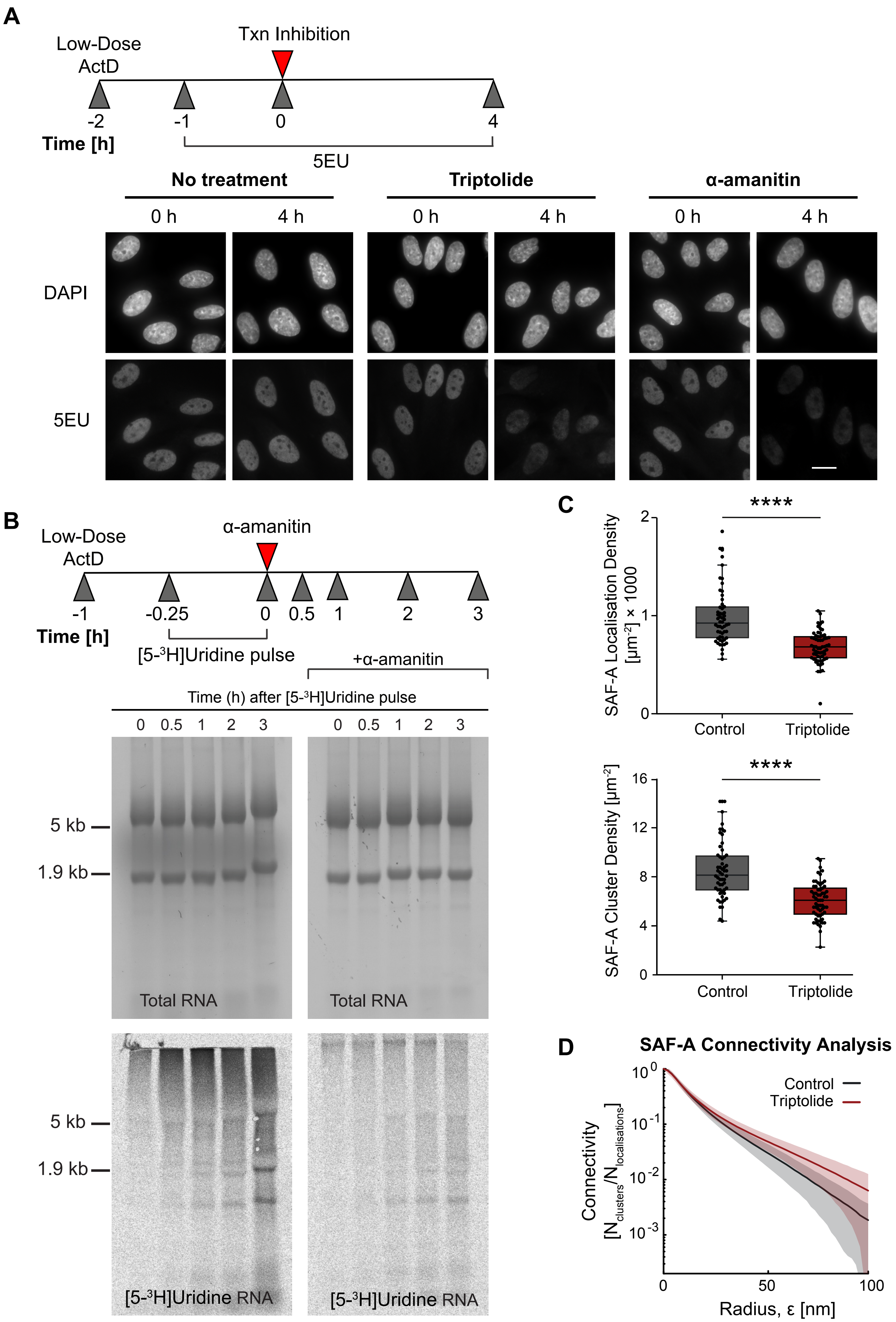


Marenda et al. Supplementary Figure 5

**A**

| | Gene.names | Protein.names |
|---|---|---|
| 1 | HNRNPU/SAF-A | Heterogeneous nuclear ribonucleoprotein U |
| 2 | HNRNPUL1 | Heterogeneous nuclear ribonucleoprotein U-like protein 1 |
| 3 | HNRNPUL2 | Heterogeneous nuclear ribonucleoprotein U-like protein 2 |
| 4 | SAFB | Scaffold attachment factor B1 |
| 5 | SAFB2 | Scaffold attachment factor B2 |
| 6 | HNRNPK | Heterogeneous nuclear ribonucleoprotein K |
| 7 | HNRNPF | Heterogeneous nuclear ribonucleoprotein F |
| 8 | HNRNPA1 | Heterogeneous nuclear ribonucleoprotein A1 |
| 9 | HNRNPDL | Heterogeneous nuclear ribonucleoprotein D-like |
| 10 | HNRNPL | Heterogeneous nuclear ribonucleoprotein L |
| 11 | HNRNPH1 | Heterogeneous nuclear ribonucleoprotein H |
| 12 | HNRNPH3 | Heterogeneous nuclear ribonucleoprotein H3 |
| 13 | HNRNPAB | Heterogeneous nuclear ribonucleoprotein A/B |
| 14 | HNRNPR | Heterogeneous nuclear ribonucleoprotein R |
| 15 | HNRNPA0 | Heterogeneous nuclear ribonucleoprotein A0 |
| 16 | HNRNPA2B1 | Heterogeneous nuclear ribonucleoproteins A2/B1 |
| 17 | HNRNPD | Heterogeneous nuclear ribonucleoprotein D0 |
| 18 | HNRNPA3 | Heterogeneous nuclear ribonucleoprotein A3 |
| 19 | SYNCRIP | Heterogeneous nuclear ribonucleoprotein Q |
| 20 | MATR3 | Matrin-3 |
| 21 | PARP1 | Poly [ADP-ribose] polymerase 1 |
| 22 | EWSR1 | RNA-binding protein EWS |
| 23 | FUS | RNA-binding protein FUS |
| 24 | TAF15 | TATA-binding protein-associated factor 2N |
| 25 | XRN2 | 5-3 exoribonuclease 2 |
| 26 | PABPC1 | Polyadenylate-binding protein 1 |
| 27 | PABPC4 | Polyadenylate-binding protein 4 |
| 28 | PABPN1 | Polyadenylate-binding protein 2 |
| 29 | CPSF4 | Cleavage and polyadenylation specificity factor subunit 4 |
| 30 | NUDT21 | Cleavage and polyadenylation specificity factor subunit 5 |
| 31 | CPSF7 | Cleavage and polyadenylation specificity factor subunit 7 |
| 32 | CPSF6 | Cleavage and polyadenylation specificity factor subunit 6 |
| 33 | RNASEH2A | Ribonuclease H2 subunit A |
| 34 | RNASEH2B | Ribonuclease H2 subunit B |
| 35 | RPA1 | Replication protein A 70 kDa DNA-binding subunit |
| 36 | RPA2 | Replication protein A 32 kDa subunit |

**B**

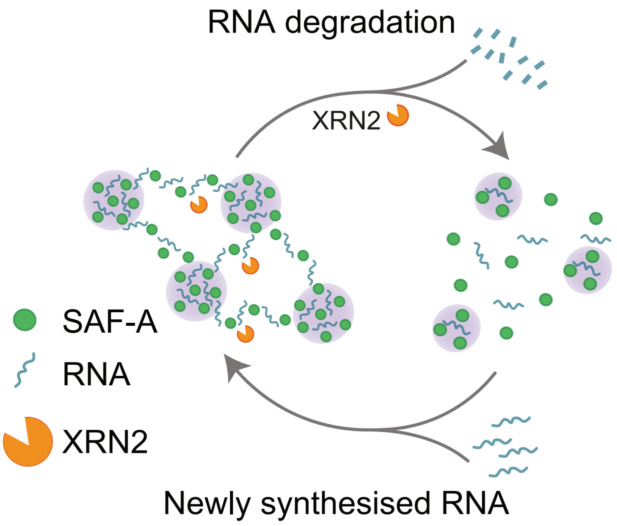


Marenda et al. Supplementary Figure 6

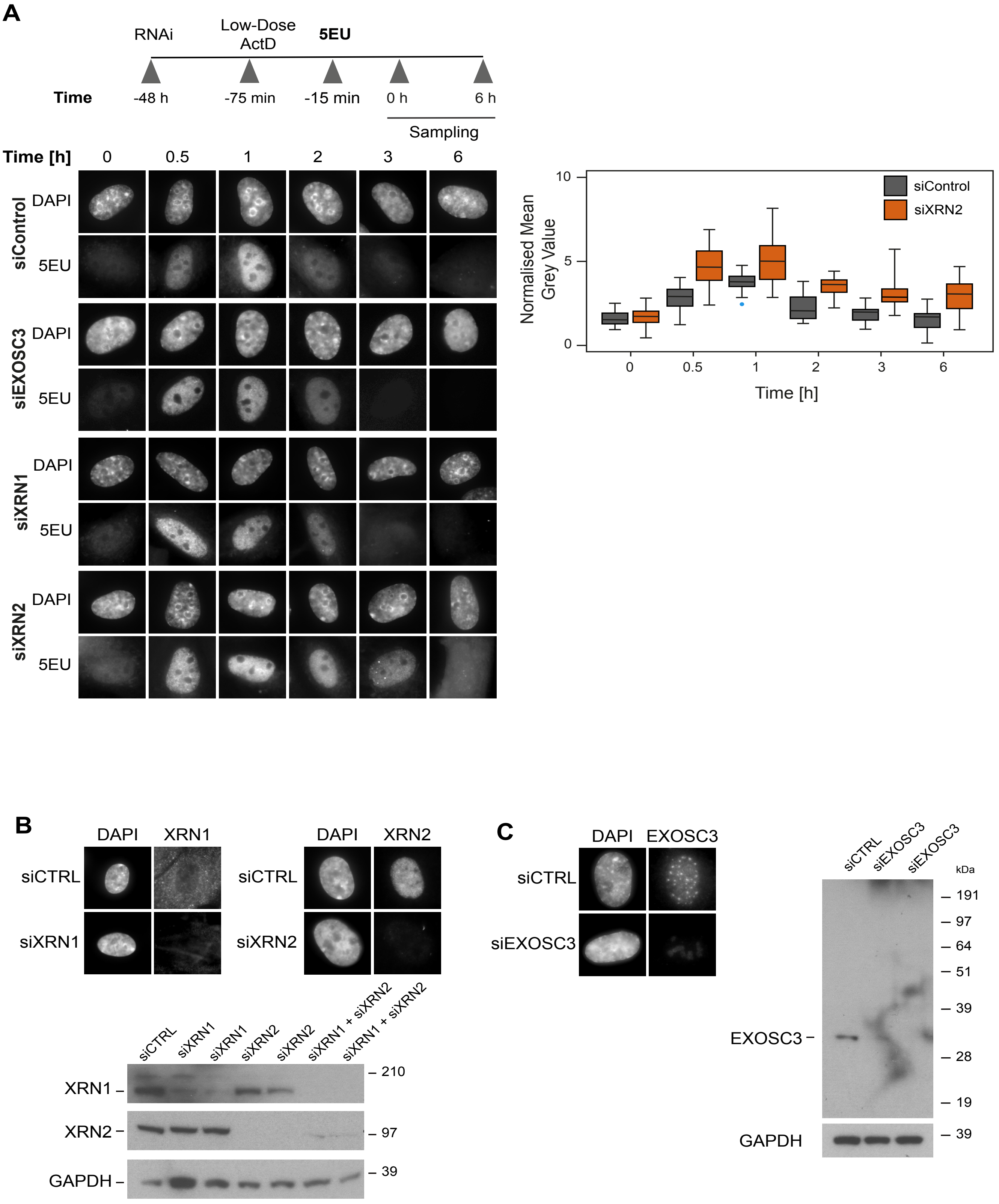


Marenda et al. Supplementary Figure 7

D

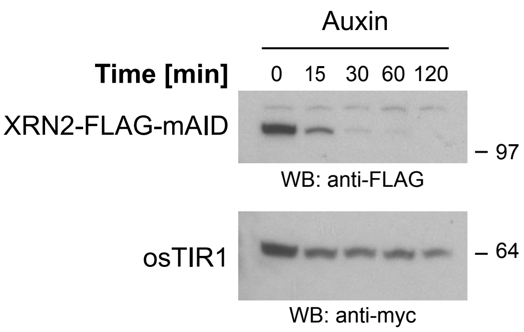


E

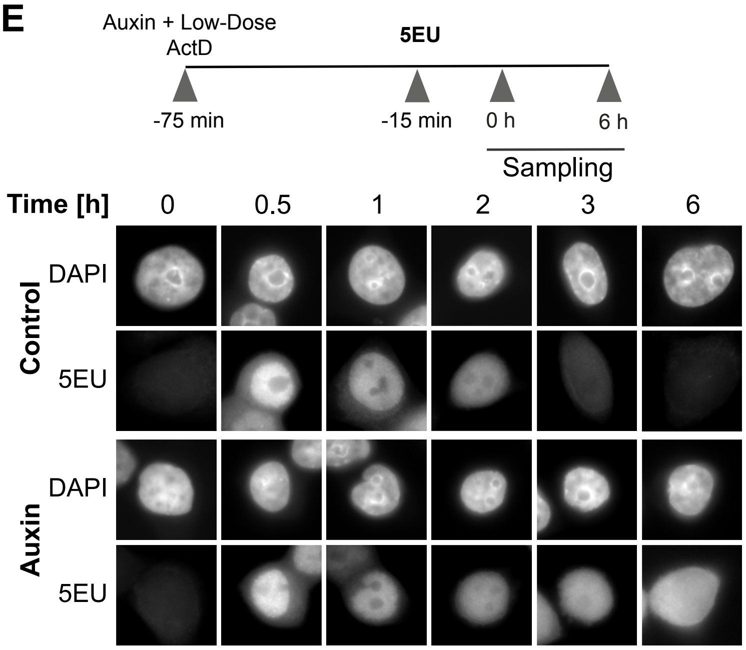


Marenda et al. Supplementary Figure 7

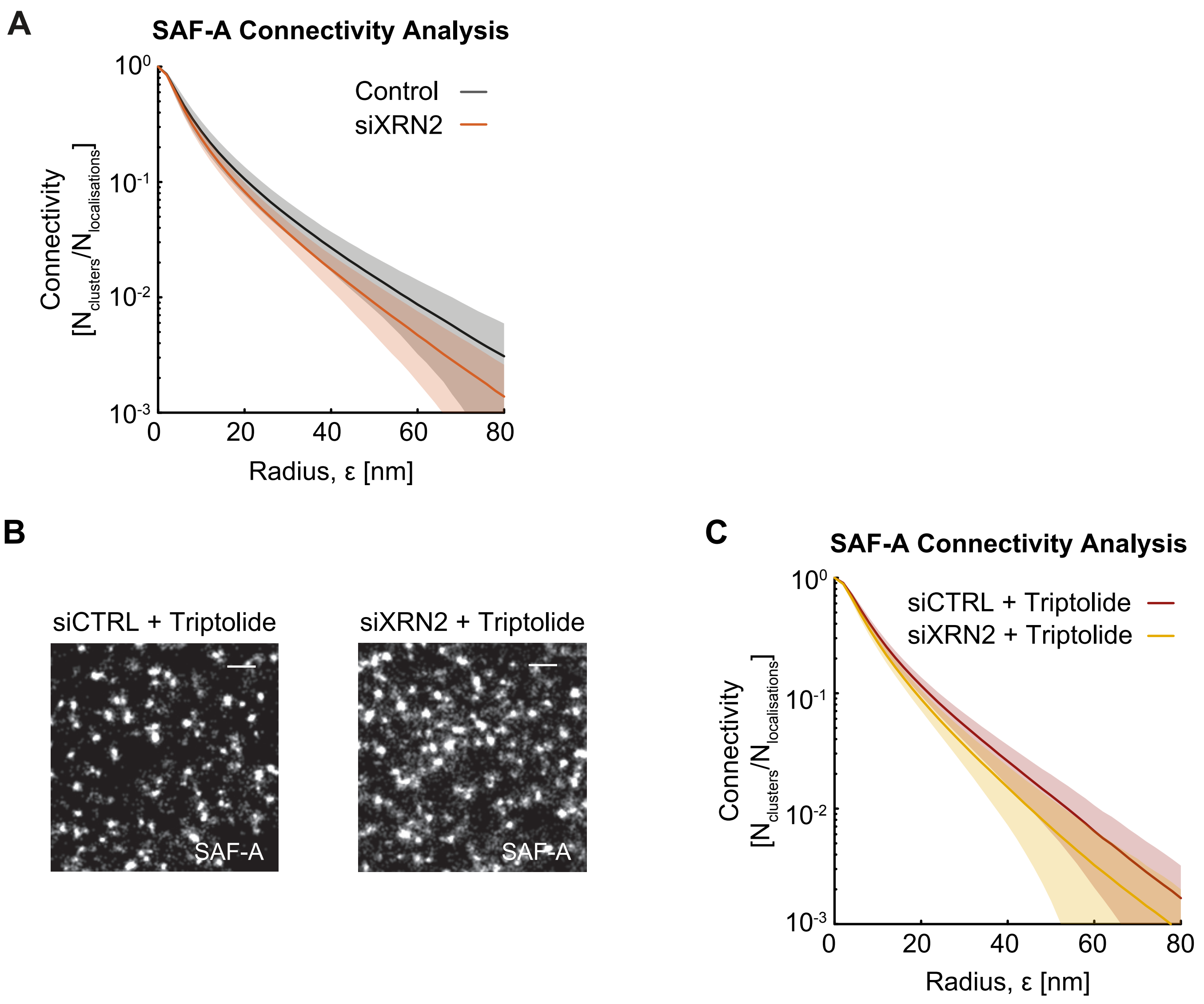


Marenda et al. Supplementary Figure 8

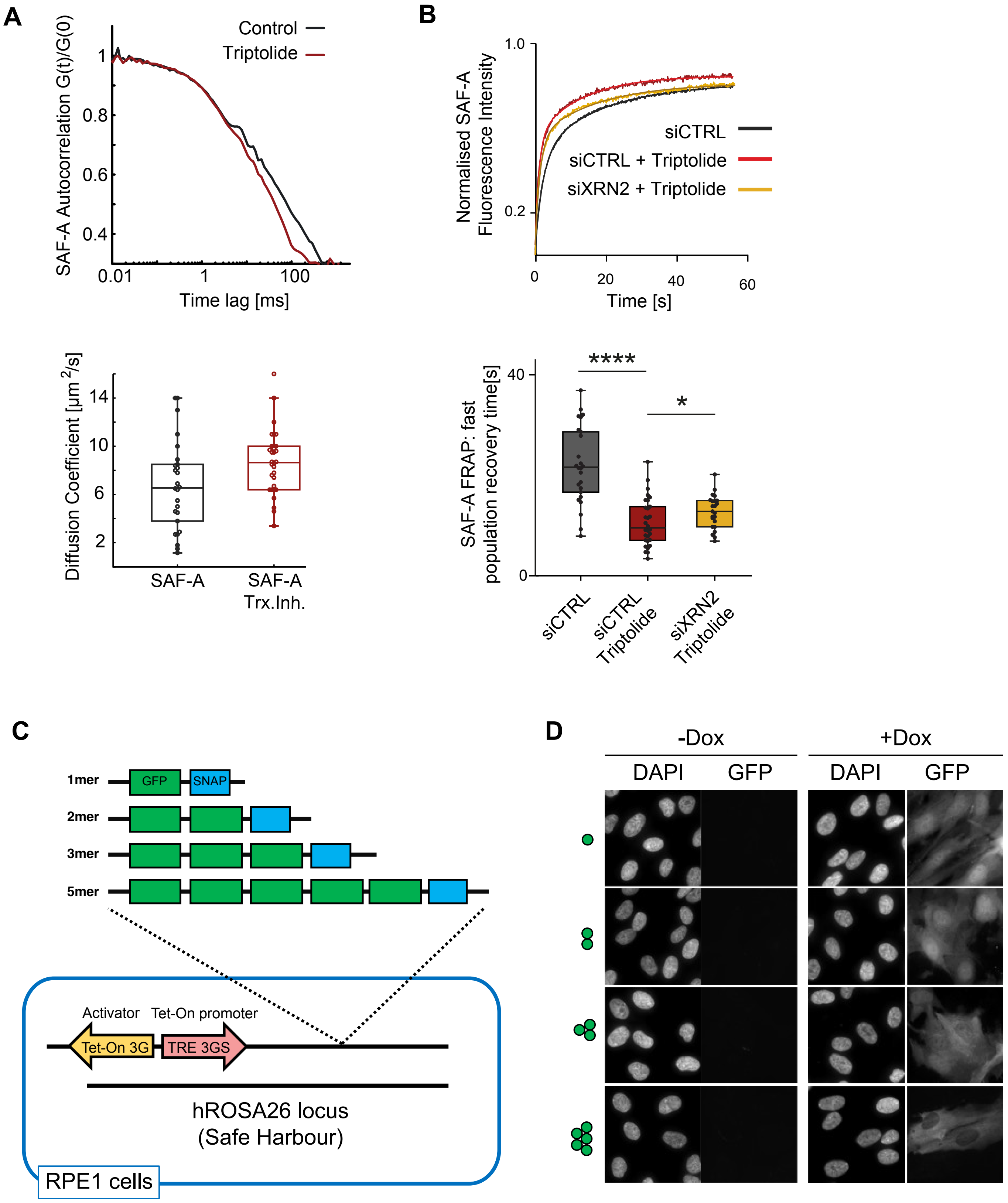


Marenda et al. Supplementary Figure 9
